# Processing and release of the maize phytocytokine Zip1

**DOI:** 10.1101/2025.06.11.659102

**Authors:** Maurice Koenig, Zarah Sorger, Parisa Kakanj, Paula Dewes, Melissa Mantz, Andreas Perrar, Muthusaravanan Sivaramakrishnan, Simon Stael, Balakumaran Chandrasekar, Pitter F. Huesgen, Johana Misas Villamil, Gunther Doehlemann

## Abstract

Phytocytokines are endogenous peptides that modulate plant immunity outcomes, yet how their maturation and spatial deployment are controlled remains unclear. Here we show that the maize phytocytokine precursor PROZIP1 is controlled by a spatially separated, two-stage proteolytic pathway that mechanistically uncouples signal activation from extracellular attenuation. PROZIP1 associates with the endoplasmic reticulum and undergoes intracellular, arginine-dependent processing by type II metacaspases, generating a C-terminal PROZIP1 fragment (Ct-PROZIP1). This processing licenses PROZIP1 for export to the apoplast via an ER-Golgi-independent route. Proteomic mapping and mutational analyses identify arginine residues flanking the Zip1 peptide as critical for efficient processing and secretion. The calcium-dependent metacaspase ZmMC9 specifically processes PROZIP1, thereby efficiently generating the bioactive Ct-PROZIP1 fragment. In the apoplast, Ct-PROZIP1 is further processed by papain-like cysteine proteases and additional extracellular proteases, contributing to Zip1 turnover and signal clearance. While the free Zip1 peptide is detected at later stages, Ct-PROZIP appears to be the primary signaling entity in modulating pathogen-induced immune responses. Together, these findings demonstrate a previously unknown complexity in peptide signaling, suggesting a multilayered control of phytocytokine activity that provides spatial and temporal precision to disease modulation in maize.

## Introduction

Plant signaling peptides, termed phytocytokines, act as secondary endogenous danger signals induced by damage and environmental stimuli and regulate growth, development and immunity. The term reflects shared conceptual features with metazoan cytokines^1–3^. However, how phytocytokines are activated and released from their precursors remains poorly understood. Wounding, pathogen infection and perception of microbe-associated molecular patterns (MAMPs) trigger phytocytokine production, and these peptides function as immunomodulators that amplify defense programs^4–6^. Phytocytokines span distinct functional classes: Several phytocytokines, including Zip1, plant elicitor peptides (Peps), PAMP-induced peptides (PIPs), serine-rich endogenous peptides (SCOOPs), GRIM REAPER peptide (GRIp) and GmSubPeps enhance extracellular immune responses and are frequently linked to salicylic acid-associated signaling^7–11^. By contrast, *CLAVATA3/EMBRYO SURROUNDING REGION–RELATED* peptides (CLEs)^12,13^ or phytosulfokines (PSKs)^14–17^ primarily regulate growth and cell fate decisions and are not associated with immune activation.

Some phytocytokines have an N-terminal signal peptide in their precursors, which determines engagement of the canonical secretory pathway^18,19^. Precursors of other phytocytokines such as Systemins, Pep1, Zip1 and GmPEPs do not have N-terminal secretion signals, which raises the question of how they reach the apoplast. Systemin is one of the mechanistically best characterized phytocytokines. Wounding induces accumulation of the active 18-amino-acid peptide, which is generated by intracellular phytaspase-mediated processing of the PROSys precursor followed by N-terminal trimming, likely by a leucine aminopeptidase^20,21^. Systemin perception by the receptor-like kinase SYR1 triggers oxidative burst, ethylene production and jasmonate pathway activation, including accumulation of 12-oxo-phytodienoic acid (OPDA) and defense gene induction^22–24^.

Proteolytic processing is generally required for phytocytokine release. In the apoplast, papain-like cysteine proteases (PLCPs) and serine proteases (subtilases) are the most abundant class of proteases^25,26^, and both participate in peptide maturation. In maize, the PLCPs CP1 and CP2 have been shown to contribute to apoplastic processing of PROZIP1, consistent with a role in extracellular phytocytokine processing and turnover rather than initial activation^7^. How PROZIP1 is initially activated and how the resulting peptide signal reaches the apoplast remain unknown. In *Arabidopsis thaliana*, Xylem Cysteine Peptidase 1 (XCP1) processes PR1 to release CAPE9^27^, and in wheat, RD21 promotes Wip1 release and resistance to wheat yellow mosaic virus^28^. Subtilases can act as processing proteases with high substrate specificity^29^, including phytaspases involved in Systemin maturation^30^, and subtilases such as SBT1.1, SBT3.8 and SBT6.1, which process PSK1, Plant Peptide Containing Sulfated Tyrosine (PSYs), CLE-like (CLEL) peptides and GOLVEN1^31–34^. Intracellular proteases also regulate phytocytokine precursors. In *A. thaliana*, Pep1 is released from PROPEP1 by the Ca²⁺-activated type II METACASPASE 4 (AtMCA-IIa, hereafter called AtMC4), which cleaves at a C-terminal arginine residue^35–37^. Similarly, in wheat TaMCA-IIa cleaves TaPROPEPs at a conserved arginine residue promoting TaPep maturation and resistance to Fusarium head blight^38^. Besides, AtMCA-IIf (hereafter called AtMC9) processes the GRI precursor at basic residues to release GRIp^11,39,40^. These findings suggest that intracellular cleavage can act as a regulatory checkpoint prior to extracellular peptide function.

Despite this progress, the subcellular localization of many phytocytokine precursors and the routes by which non-secreted peptides reach the apoplast remain poorly understood. This also holds true for Zip1, a maize-specific phytocytokine that enhances salicylic acid-mediated immunity and activates PLCPs, thereby promoting resistance to the biotrophic fungus *Ustilago maydis*^7,41^.

Here, we investigated the localization, maturation and transport to the apoplast of PROZIP1, the precursor of Zip1. We found intracellular cleavage being a prerequisite for release of a bioactive C-terminal precursor Ct-PROZIP1 that is exported to the apoplast. Ct-PROZIP1 exerts stronger effects in plant-microbe interactions than Zip1 alone, whereas free Zip1 remains bioactive but likely reflects further proteolytic processing. Notably, proteolysis also occurs within the Zip1 sequence and suggest that Ct-PROZIP1 rather than free Zip1 represents the dominant signaling form. Based on our results, we propose a two-step regulatory mechanism in which intracellular processing licenses secretion of a C-terminal Zip1-containing fragment, while subsequent apoplastic proteolysis mediates signal attenuation and peptide clearance.

## Results

### Subcellular localization and intracellular processing of PROZIP1

To examine the subcellular localization of PROZIP1, we expressed C-terminally fluorescently tagged PROZIP1 in maize protoplasts. PROZIP1 fluorescence was detected intracellularly, with signal at the nuclear periphery and in the cytoplasm, whereas NLS-mCherry was confined to the nucleus (Fig. S1A). Similar fluorescence patterns were observed in *Nicotiana benthamiana* epidermal cells expressing PROZIP1-mCherry (Fig. S1B), indicating that the fluorescent tag does not markedly alter intracellular distribution.

As PROZIP1 fluorescence accumulated at the nuclear periphery, we examined its subcellular distribution relative to the endoplasmic reticulum (ER) and compared PROZIP1 with the cleavage-site mutant PROZIP1^CS^, in which double arginine residues flanking the Zip1 region were substituted with alanine^7^. PROZIP1-GFP partially co-localized with the ER marker mCherry-KDEL^42,43^ (Fig. 1A). GFP signal overlapped with the ER marker at the nuclear periphery, but was not restricted to KDEL-positive structures, and additional cytosolic signal was detected. GFP fluorescence was also observed in the nucleolar region but was not detected for PROZIP1^CS^-GFP, which showed increased overlap with the ER marker. In this case, the GFP signal largely followed the reticulate ER pattern, although complete ER confinement was not observed. Comparable fluorescence patterns were observed in *N. benthamiana* epidermal cells (Fig. S1C). Together, these data indicate that PROZIP1 is intracellularly distributed and partially associated with the ER.

**Figure 1.**
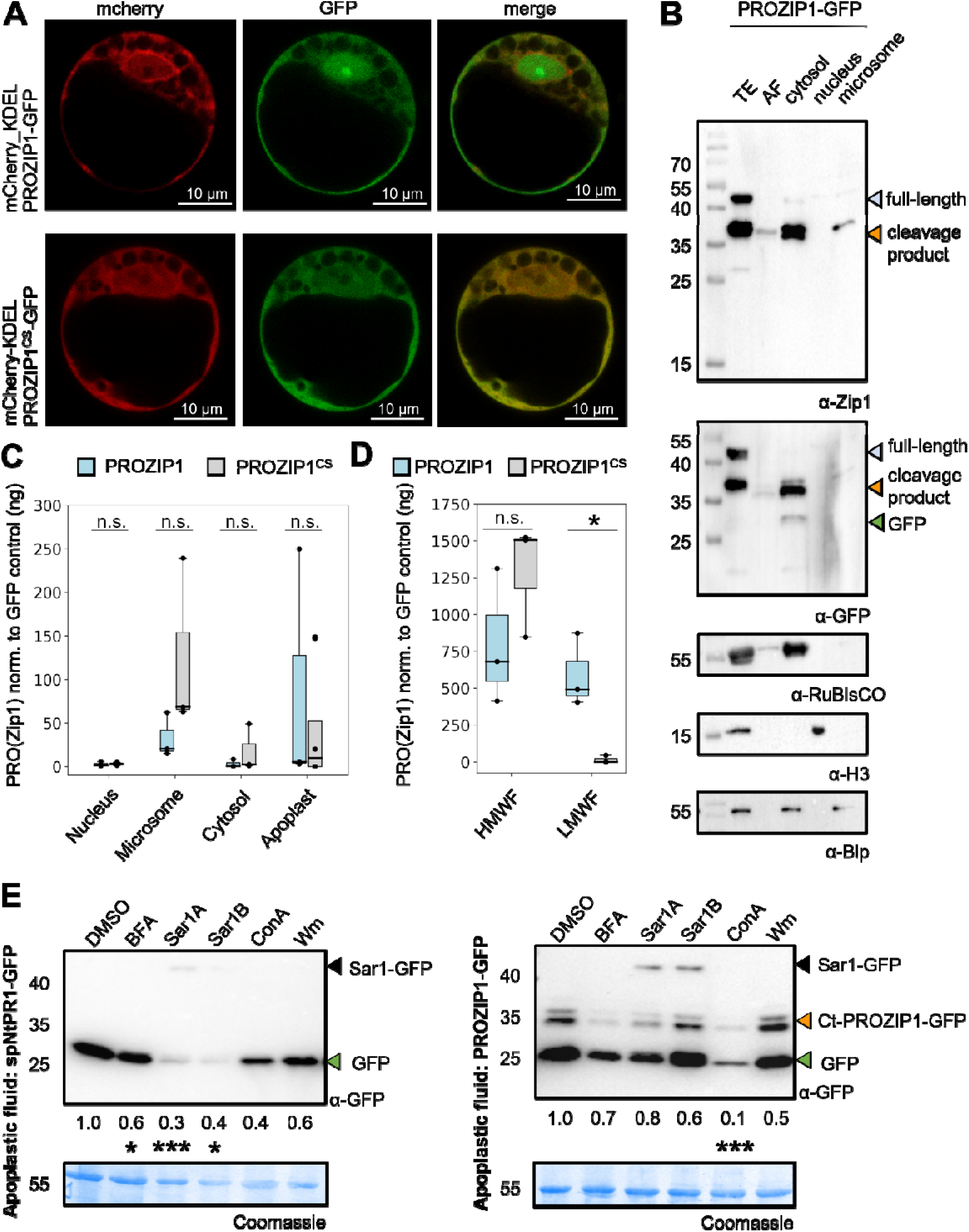
N-terminal processing of PROZIP1 enables C-terminal translocation to the apoplast. **(A)** Maize protoplasts expressing PROZIP1-GFP or the cleavage-site mutant PROZIP1^CS^-GFP together with the ER marker mCherry-KDEL. GFP (green), mCherry (red) and merged signals are shown. Both constructs display intracellular fluorescence with perinuclear enrichment and ER-associated structures. Images were acquired 16h after transfection. Scale bars, 10 μm. **(B)** Subcellular fractionation of *Nicotiana benthamiana* leaves expressing PROZIP1-GFP. Cytosolic, nuclear, microsomal and apoplastic fractions were analyzed by immunoblotting. Fraction purity was verified using RuBisCO (cytosol), histone H3 (nucleus) and BiP (microsomes). Full-length PROZIP1-GFP (blue), a cleavage product (orange) and GFP (green) are indicated. The cleavage product is detected in cytosolic, microsomal and apoplastic fractions, consistent with intracellular processing of PROZIP1 followed by an extracellular release of the processed fragment. **(C)** Quantification of Zip1-containing fragments in the fractions shown in B using an α-Zip1 ELISA. Signals from PROZIP1-GFP (blue) and PROZIP1^CS^-GFP (grey) were normalized to GFP background. Boxplots represent independent biological replicates; dots indicate individual data points. Zip1-containing fragments are detected in intracellular fractions and the apoplast. PROZIP1-derived fragments tend to accumulate in the apoplast, whereas signals from the cleavage-site mutant remain primarily associated with cytosolic and microsomal fractions. **(D)** Apoplastic fluid was separated into >5_kDa (HMWF) and <5_kDa (LMWF) fractions and analyzed by α-Zip1 ELISA. Signals were normalized to GFP background. Boxplots represent independent biological replicates. Zip1-containing fragments are enriched in the low-molecular-weight fraction in PROZIP1 samples, but are strongly reduced in the PROZIP1^CS^. **(E)** Extracellular accumulation of spNtPR1-GFP (marker for conventional secretion) and PROZIP1-GFP in *Nicotiana benthamiana* following treatment with BFA, ConA or Wm, or co-expression with dominant-negative Sar1A or Sar1B. GFP and the PROZIP1 cleavage product (Ct-PROZIP1) were detected by anti-GFP immunoblotting. Band intensities were quantified; numbers below the lanes indicate values normalized to DMSO. Ct-PROZIP1 accumulation in the apoplast is insensitive to BFA or Sar1 but reduced by ConA. Statistics: one-tailed paired Student’s t-test (P < 0.05; n.s.).

To analyze PROZIP1 processing, *N. benthamiana* leaves expressing PROZIP1-GFP were subjected to subcellular fractionation followed by immunoblotting using antibodies against GFP and Zip1, as well as the compartment markers RuBisCO (cytosol), histone H3 (nucleus) and BiP (microsomes). Total extracts contained full-length PROZIP1-GFP (∼43□kDa) and multiple Zip1-containing fragments (∼28-36□kDa), whereas PROZIP1^CS^-GFP accumulated predominantly as full-length protein with altered cleavage products (Fig. 1B; Fig. S2A). Intracellularly processed PROZIP1-GFP fragments were detected in cytosolic and microsomal fractions but not in nuclear fractions, consistent with the ER-proximal localization inferred from mCherry-KDEL colocalization. By contrast, PROZIP1^CS^-GFP showed reduced microsomal processing and increased stability of the full-length protein, indicating altered intracellular cleavage (Fig. S2A). In apoplastic fractions, PROZIP1-GFP accumulated predominantly as a ∼36□kDa Zip1-containing fragment, matching the cytosolic cleavage product and indicating export of a processed C-terminal fragment (Ct-PROZIP1) from the cytosol to the apoplast (Fig. 1B). By contrast, PROZIP1^CS^-GFP was detected mainly as full-length protein, indicating strongly reduced accumulation of processed PROZIP1 species in the apoplast (Fig. S2A). These data suggest that PROZIP1 undergoes intracellular N-terminal processing at double-arginine (RR) sites prior to secretion.

To test whether the N-terminal region undergoes site-specific cleavage, GFP fusions of PROZIP1^1-70^ and PROZIP1^CS_1-70^ were analyzed via western blot (Fig. S2B). PROZIP1^1-70^-GFP displayed discrete cleavage products in cytosolic and microsomal fractions, whereas the mutant showed pronounced laddering, consistent with aberrant processing or destabilization. A ∼20□kDa fragment was also present in GFP controls, indicating GFP-derived cleavage. Hydropathy analysis revealed a decreasing hydrophobicity gradient from the N- to the C-terminus, with the N-terminal region being relatively more lipophilic, consistent with transient membrane association rather than stable insertion (Fig. S2C).

To quantify Zip1-containing fragments, subcellular fractions were analyzed by ELISA using a Zip1-specific antibody^44^. Zip1-containing species were detected in cytosolic, microsomal and apoplastic fractions for both constructs, with PROZIP1-GFP displaying relatively lower intracellular levels and higher apoplastic levels compared with the cleavage-site mutant (Fig. 1C). Size fractionation of apoplastic fluids showed that Zip1 was present exclusively in the low-molecular-weight fraction of PROZIP1-GFP samples, whereas release of free Zip1 was significantly reduced in PROZIP1^CS^-GFP (Fig. 1D), confirming impaired peptide liberation.

To test by which route Ct-PROZIP1 is secreted, PROZIP1-GFP was co-expressed with Sar1A or Sar1B, or leaves were treated with brefeldin A (BFA), concanamycin A (ConA) or wortmannin (Wm). Sar1 overexpression and BFA inhibited secretion of the conventional marker spNtPR1-GFP^45^ (Fig. 1E, left panel), consistent with blockade of ER-Golgi trafficking^46–49^. In contrast, Ct-PROZIP1-GFP accumulation in the apoplast was unaffected by BFA or Sar1, but it was significantly reduced by ConA, a V-ATPase inhibitor that disrupts vacuolar and endomembrane trafficking^50,51^ (Fig. 1E, right panel). These results indicate that Ct-PROZIP1 export likely occurs via a Golgi-independent secretion route that requires compartment acidification, rather than through the canonical ER-Golgi pathway, consistent with unconventional protein secretion^52^.

### Intracellular arginine-dependent processing of PROZIP1

To map *in vivo* cleavage sites, we applied amino-terminal oriented mass spectrometry of substrates (ATOMS)^53^ to PROZIP1-GFP and PROZIP1^CS^-GFP expressed in both maize protoplasts and *N. benthamiana* leaves. GFP-tagged C-terminal fragments were enriched by immunoprecipitation, and neo-N termini were labelled by reductive dimethylation with ^13^CDO₂-formaldehyde. Mass spectrometry predominantly identified C-terminal cleavage products, consistent with selective enrichment of GFP-tagged fragments. Identified peptides were highly similar between maize and *N. benthamiana*, indicating conserved processing across species (Fig. 2A; Fig. S3).

**Figure 2.**
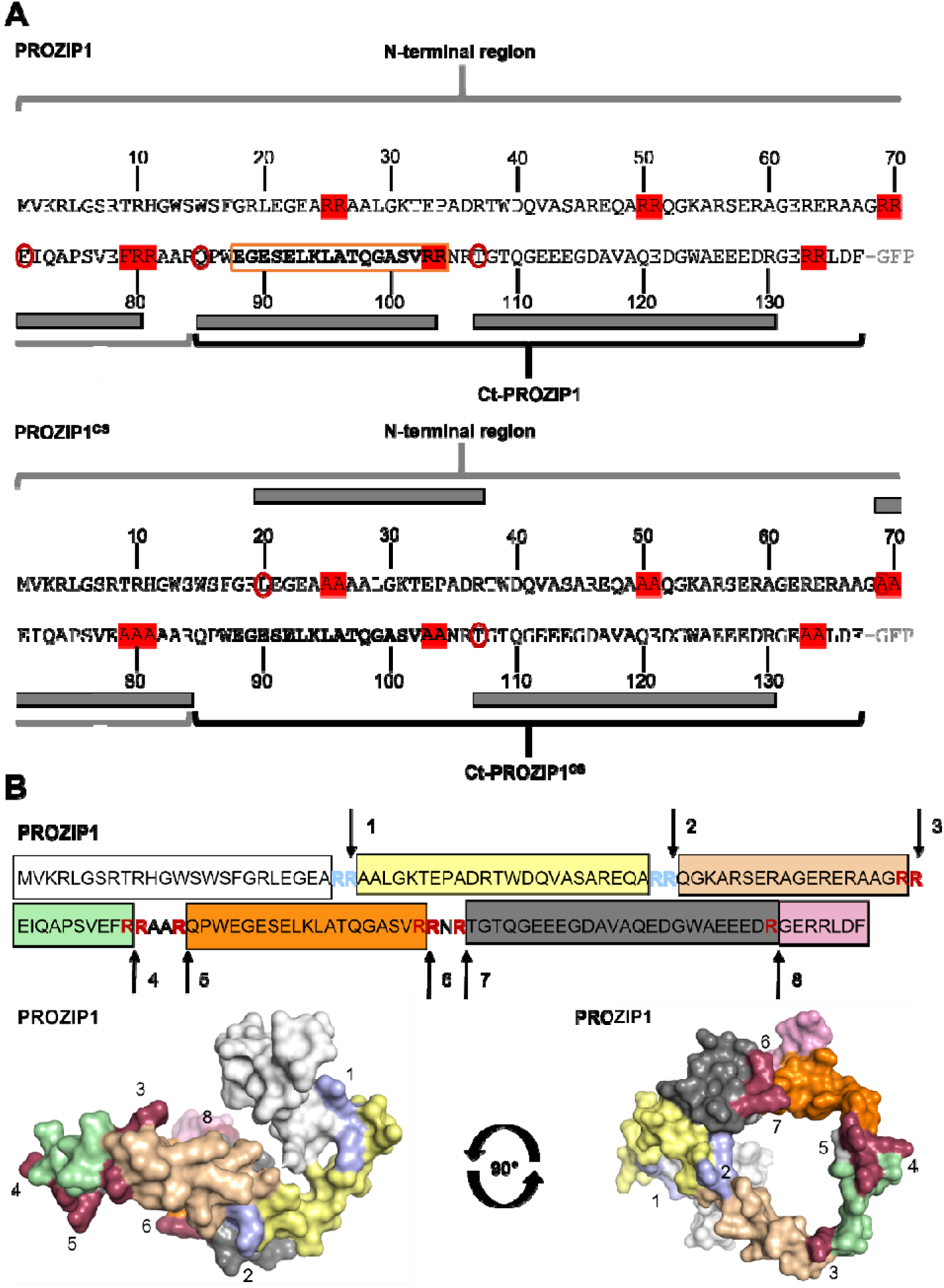
Arginine-dependent processing of PROZIP1 is. **(A)** Identification of PROZIP1 cleavage sites by N-terminomics. PROZIP1-GFP or PROZIP1^CS^-GFP were expressed in maize protoplast and enriched by pull-down. Newly generated N termini were labelled by reductive dimethylation and identified by mass spectrometry. Detected neo-N termini are indicated as red circles, and corresponding peptides are mapped as grey boxes onto the sequences. The Zip1 peptide is highlighted in orange, and Arg-to-Ala substitutions in PROZIP1^CS^ are shown in red. Neo-N termini were detected immediately downstream of RR□□^-^□□, R□□ and R¹□□, indicating cleavage after arginine residues. Cleavage at R□□ and R¹□□ flanks the Zip1 region and is consistent with release of the Zip1-containing fragment, which is absent in PROZIP1^CS^. **(B)** Predicted three-dimensional structure of PROZIP1. The PROZIP1 sequence (top) and surface models of PROZIP1 (bottom) are shown in two orientations. Peptides identified by mass spectrometry are highlighted in the sequence and models. The Zip1 region is shown in orange; putative cleavage sites inferred from immunoblotting are shown in light blue, and cleavage sites identified by mass spectrometry in dark red. Arrows indicate cleavage positions, numbered from the N to the C terminus, and corresponding sites are marked in the models. The identified cleavage sites are surface-exposed in the model, consistent with protease accessibility.

Cleavage sites were arginine-dependent, with most neo-N termini adjacent to paired arginine residues. The peptide EIQAOSVEFR^71-80^, immediately upstream of Zip1, was extended in PROZIP1^CS^, indicating altered processing upon mutation of neighboring RR sites (RR^69-70^ and RR^79-81^) (Fig. 2A). A peptide overlapping the previously defined Zip1 sequence (EGESELKLATQGASVRR^88-104^)^7^ and detection of a downstream neo-N terminus at T^107^ support cleavage both upstream and downstream of Zip1, consistent with peptide release *in vivo*. In line with this, the dominant ∼36□kDa Ct-PROZIP1^85-137^-GFP fragment observed by immunoblotting (Fig. 1B) likely results from cleavage at upstream RR sites (RR^25-26^, RR^50-51^, RR^69-70^, RR^80-81^), which generates Q^85^ as the neo-N terminus. No free Zip1 peptide was detected in PROZIP1^CS^-GFP (Fig. 2A), indicating that RR mutations impair Zip1 release and C-terminal translocation. Additional cleavage at single arginine residues flanking Zip1 (R^84^, R^106^) suggests that monobasic sites can also contribute to Zip1 maturation.

Structural prediction of PROZIP1 using RoseTTAFold suggests a compact, ring-like architecture in which the N-terminal region, the central Zip1 peptide region and the acidic C-terminal region are arranged in close spatial proximity (Fig. 2B). In this model, several arginine residues that were experimentally identified as cleavage sites are surface-exposed, including sites R^26^, R^84^, R^103^ and R^106^. These sites are located on peripheral regions of the fold, consistent with their susceptibility to proteolytic processing. Together, the model supports an arginine-dependent cleavage mechanism governed by structural accessibility of P1 residues. Substitutions of these arginine residues by alanine abolish the corresponding arginine-dependent cleavage sites thereby preventing processing at these sites.

### Type II metacaspases mediate arginine-dependent processing of PROZIP1

The intracellular arginine-dependent cleavage pattern of PROZIP1 implicates metacaspases (MCAs), which preferentially cleave after basic residues and process phytocytokine precursors in *Arabidopsis thaliana*. AtMC4 cleaves PROPEP1 and AtMC9 processes GRI, releasing Pep1 and GRIp, respectively^11,36^. Both proteases are type II MCAs characterized by an extended linker between the p20 and p10 regions and the absence of an N-terminal prodomain^54^ (Fig. S4A). We initially examined the ability of recombinant AtMC4 to process PROZIP1 *in vitro*. AtMC4 cleaved PROZIP1 but not the cleavage-site mutant PROZIP1^CS^ (Fig. S4B, left panel). Notably, Zip1 abundance was unaffected, indicating that cleavage occurs outside the Zip1 peptide region (Fig. S4B, right panel). This prompted us to extend our analysis to maize MCAs.

Phylogenetic analysis identified eight type I and three type II maize MCAs, with *ZmMC9* (ZmMCA-IIa) and *ZmMC10* (ZmMCA-IIb) clustering with the Ca^2+^-dependent *AtMC4*, and *ZmMC11* (ZmMCA-IIc) grouping with the low-pH-dependent *AtMC9* (Fig. 3A). While catalytic residues are conserved, Ca²⁺-binding features and overall sequence similarity differ among type II metacaspases^54,55^. Notably, *AtMC9* and *ZmMC11* show reduced conservation and lower overall similarity, indicating functional divergence, whereas *ZmMC9* and *ZmMC10* retain greater similarity to the Ca²⁺-dependent *AtMC4*, supporting their candidacy as proteases involved in PROZIP1 processing (Fig. S4C). Further, *ZmMC9* was constitutively expressed under mock, salicylic acid, and Zip1 conditions and was the only metacaspase detected at moderate transcript abundance in our RNA-seq dataset, which was analyzed at the 6 h time point corresponding to the peak of *Prozip1* induction upon salicylic acid and Zip1 treatment (Fig. 3B; S4D). By contrast, *ZmMC10* and *ZmMC11* were expressed at near-background levels (∼1 TPM), thereby prioritizing ZmMC9 as the most plausible protease candidate during immune activation (Fig. 3B). To explore whether the maize type II metacaspase ZmMC9 can directly engage PROZIP1 as a substrate, we generated a structural docking model of the ZmMC9 p20-p10 heterodimer in complex with PROZIP1. In the resulting model, PROZIP1 associates with the metacaspase surface such that the Zip1-containing region is oriented toward the active-site cleft formed between the p20 and p10 regions (Fig. S4E). Notably, the previously proposed cleavage site at R^84^ is positioned in close proximity to the catalytic His^88^-Cys^141^ dyad of ZmMC9 (Fig. S4F), consistent with a potential protease-substrate interaction. The model supports the hypothesis that ZmMC9 can engage PROZIP1 at this site. Based on this structural prediction, we next experimentally tested whether ZmMC9 mediates cleavage of PROZIP1 at R^84^ generating Ct-PROZIP1.

**Figure 3.**
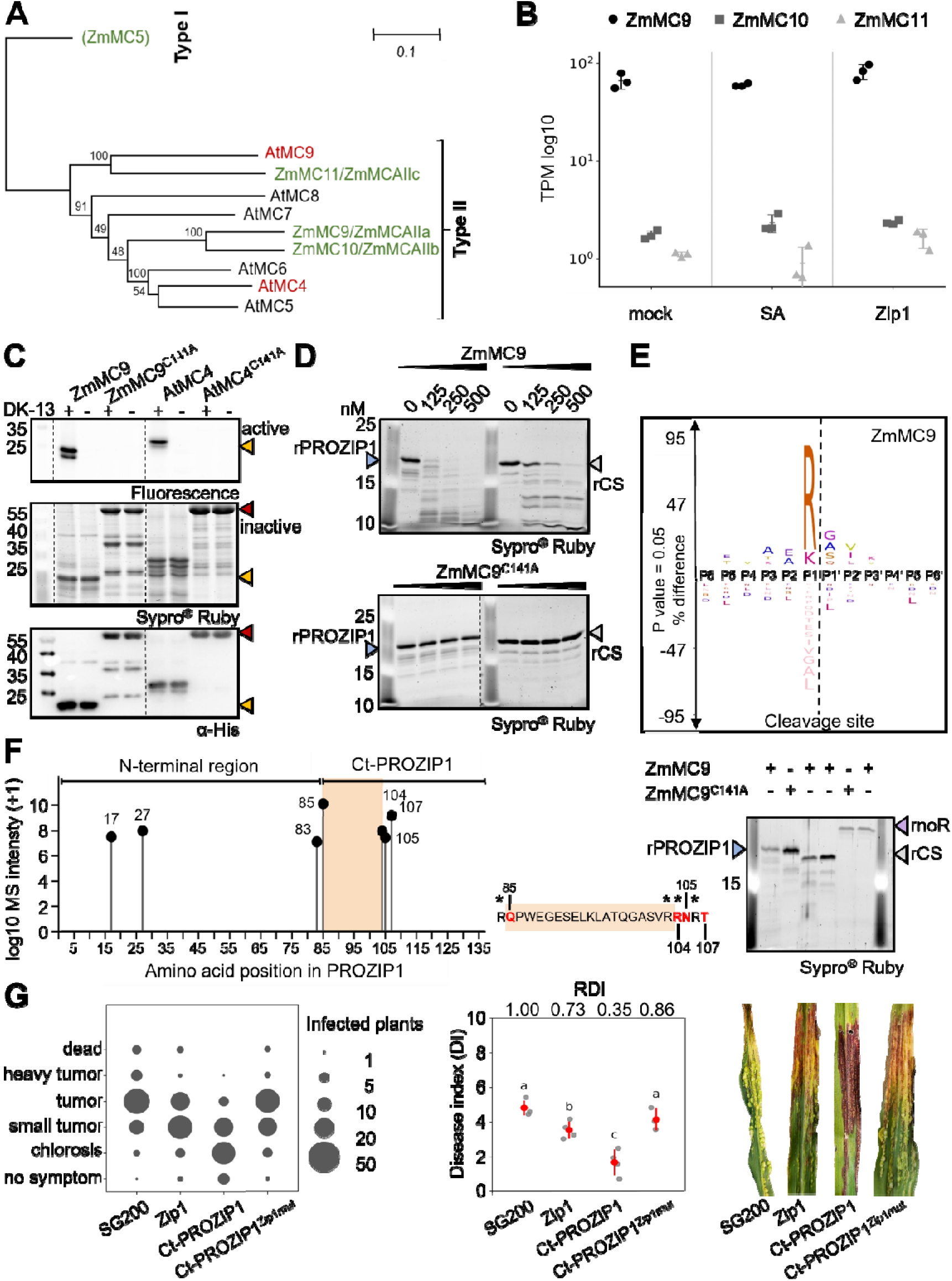
Type II metacaspase ZmMC9 processes PROZIP1 to generate Ct-PROZIP1. **(A)** Phylogenetic analysis of selected *Arabidopsis thaliana* and *Zea mays* metacaspases based on catalytic domain sequences adapted from Luan *et al.* (2021). Bootstrap values are indicated at nodes. ZmMC9 clusters within the type II metacaspase clade together with Arabidopsis homologues. Scale bar indicates substitutions per site. **(B)** RNA seq analysis of mock-, SA- and Zip1-treated maize leaves shows differential expression of three type II metacaspases (ZmMC9, ZmMC10, ZmMC11). Transcript abundance is shown as transcripts per million (TPM, log10). ZmMC9 shows significantly higher expression levels under mock and stress conditions compared to ZmMC10 and ZmMC11. **(C)** Activity-based probe labelling of metacaspases using the fluorescent probe DK-13. Recombinant ZmMC9 and AtMC4 show probe labelling, whereas catalytically inactive variants (ZmMC9^C141A^ and AtMC4^C141A^) do not. Corresponding Sypro® Ruby staining and anti-His immunoblot confirm protein loading. Arrowheads indicate labelled active protease species (yellow) and full-length proteins (red). **(D)** *In vitro* cleavage assay of PROZIP1 and PROZIP1^CS^ by increasing concentrations of ZmMC9. Reactions were incubated for 30 min and analyzed by Sypro® Ruby staining. Cleavage products accumulate in PROZIP1 (blue) but are strongly reduced in PROZIP1^CS^ (grey) samples. ZmMC9^C141A^ was used as inactive control. **(E)** Positional amino acid preferences of ZmMC9 determined by Proteomic Identification of protease Cleavage Sites (PICS). Logo plot depicts enriched residues surrounding the cleavage site (P6 to P6′), with statistically significant positions indicated. ZmMC9 shows a strong preference for arginine at the P1 position. **(F)** Mapping of ZmMC9-dependent cleavage sites in PROZIP1 by N-terminomics. Detected neo-N termini are plotted along the PROZIP1 sequence; lollipop height corresponds to log_₁₀_-transformed MS1 intensity (+1). The N-terminal region (aa 1–84), Ct-PROZIP1 (aa 85–137) and the Zip1 peptide region (aa 85–104) are indicated. The PROZIP1 sequence is shown below, with residues corresponding to identified neo-N termini highlighted in red. ZmMC9-dependent cleavage sites flank the Zip1 region, consistent with proteolytic release of Ct-PROZIP1. Mutation of the corresponding arginine residues strongly reduces (PROZIP1^CS^) or abolishes (PROZIP1^noR^) cleavage *in vitro*. **(G)** Zip1, Ct-PROZIP1 and the C-terminal region of PROZIP1^CS^ were delivered to the maize apoplast using the Trojan horse approach (van der Linde *et al*., 2018). Leaf-whorl infections were performed on 1-week-old maize plants, with strain SG200 as control. Disease symptoms were scored at 8_days post-infection (dpi), and absolute (DI) and relative disease indices (RDI) were calculated and plotted. Representative images of infected plants are depicted. Grey dots represent biological replicates; error bars indicate mean_±_s.e.m. Statistical significance was assessed by one-way ANOVA followed by Tukey’s HSD test. Ct-PROZIP1 reduces disease symptoms more strongly than Zip1, and mutation of the Zip1 motif abolishes activity, indicating that an intact Zip1 motif is required.

Recombinant ZmMC9 and active site mutant ZmMC9^C141A^ were purified from *E. coli* (Fig. S5A), and protease activity was assessed by activity-based protein profiling (ABPP) using the fluorescent probe DK-13^56^ (Fig. S5B). Active-site mutants of AtMC4 and ZmMC9 (AtMC4^C141A^, ZmMC9 ^C141A^) lacked autocatalytic processing and were not labeled by the probe confirming their inactivity (Fig. 3C; ^36^). ZmMC9 activity is Ca²⁺ dependent (Fig. S5C) and it was inhibited by leupeptin and EDTA but not by E-64 (Fig. S5D), consistent with a metacaspase inhibitory profile. To analyze PROZIP1 cleavage by ZmMC9, PROZIP1, PROZIP1^CS^ and an arginine-free variant of the pro-peptide (PROZIP1^noR^) were purified from *E. coli* (Fig. S6A-I) and subjected to *in vitro* cleavage assays. ZmMC9 processed PROZIP1 under the tested conditions (Fig. 3D; S6J). Efficient processing of PROZIP1 occurred with 250□nM ZmMC9 within 30□min, while PROZIP1^CS^ remained largely intact (Fig. 3D).

Proteomic identification of protease cleavage sites (PICS)^57^, a peptide-library-based method to determine protease substrate specificity, revealed a strong preference of ZmMC9 for arginine at the P1 position, with limited sequence constraints at adjacent positions, consistent with the substrate specificity previously described for AtMC4 and AtMC9^55,58^ and aligns with the differential processing observed for PROZIP1 and PROZIP1^CS^ *in vivo* (Fig. 1B; Fig. S2A-B) and *in vitro* (Fig. 3E).

To assess whether ZmMC9 generates Ct-PROZIP1, we co-incubated PROZIP1, PROZIP1^CS^ and PROZIP1^noR^ with ZmMC9 for subsequent analysis by ATOMS (Fig. 3F; S7A-C). ZmMC9^C141A^ served as a background control. ZmMC9-dependent cleavage occurred at the N-terminus (RR^25-26^↓A^27^) and at sites flanking Zip1 (R^84^↓Q^85^ and R^106^↓T107). Cleavage at R^84^↓Q^85^ is consistent with the generation of Ct-PROZIP1. Inactive ZmMC9 did not cleave at R^84^↓Q^85^ and consequently did not produce Ct-PROZIP1 (Fig. S7B). Compared with wild-type PROZIP1, PROZIP1^CS^ and PROZIP1^noR^ showed impaired processing, with reduced or absent productive cleavage and heterogeneous neo-N termini within the Zip1 sequence, consistent with non-productive peptide degradation (Fig. 3F, right panel; Fig. S7A and S7C).

We next tested whether Ct-PROZIP1 is biologically active or whether proteolytic release of Zip1 is required for signaling. To address this, we used the Trojan horse delivery system to introduce Zip1 or Ct-PROZIP1 into the maize apoplast via the biotrophic fungus *U. maydis*^7,59^. Whereas Zip1 reduced disease severity relative to the pathogenic control strain, delivery of Ct-PROZIP1 caused a markedly stronger phenotype, with more reduced tumor formation and pronounced chlorosis (Fig. 3G; Fig. S8). These results indicate that Ct-PROZIP1 itself is a bioactive signaling peptide and suggest that the C-terminal precursor rather than free Zip1 represents the dominant functional form in the apoplast. Removal of the Zip1 motif from Ct-PROZIP1 (Ct-PROZIP1^Zip1mut^) abolished this effect, indicating that the Zip1 sequence is required for Ct-PROZIP1 bioactivity in the apoplast and that further proteolysis primarily contributes to signal attenuation rather than peptide maturation.

### Apoplastic PLCPs cleave C-terminal PROZIP1

Previous work suggested that the PLCPs CP1 and CP2 contribute to Zip1 release from PROZIP1 in the apoplast^7^. The finding of an intracellular, metacaspase-dependent processing step therefore raised the question of whether PLCPs might also participate in the intracellular cleavage of PROZIP1, or whether they only act on the secreted C-terminal fragment.

To test whether PLCPs contribute to intracellular processing, maize protoplasts expressing PROZIP1-GFP or PROZIP1^CS^-GFP were treated with the cell-permeable PLCP inhibitor E64d (Fig. 4A-B). Activity-based protein profiling using the biotinylated probe DCG04 showed efficient inhibition of intracellular PLCP activity under these conditions (Fig. 4A)^60^. However, the E64d treatment did not alter the subcellular localization or cleavage patterns of either construct (Fig. 4A, 4B), indicating that PLCP activity is not required for the intracellular processing of PROZIP1.

**Figure 4.**
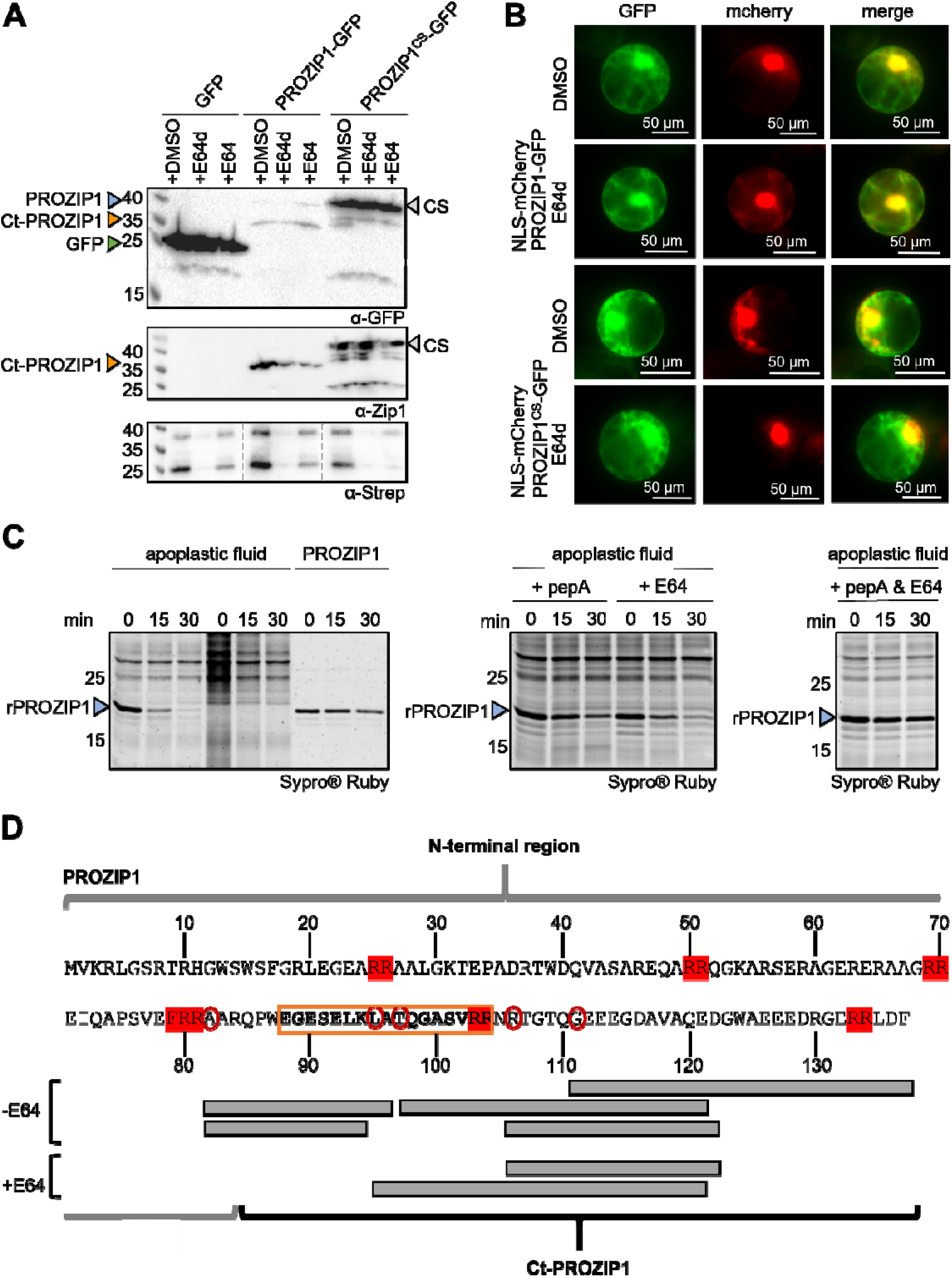
Apoplastic PLCPs contribute to release and clearance of Zip1. **(A)** Protein extracts of maize protoplast expressing PROZIP1-GFP and PROZIP1^CS^-GFP were analyzed by immunoblotting using α-Zip1 and α-GFP antibodies. PLCP activity was assessed by ABPP using DCG-04-biotin and detected by streptavidin immunoblotting. The cleavage pattern of PROZIP1 and PROZIP1^CS^ is unchanged, indicating that intracellular PLCPs are not required for PROZIP1 processing. **(B)** PROZIP1-GFP and PROZIP1^CS^-GFP were co-expressed with NLS-mCherry in maize protoplasts and treated with the cell-permeable PLCP inhibitor E64d or DMSO. Subcellular localization was analyzed by fluorescence microscopy 16_h after transfection. PLCP inhibition does not alter intracellular localization of PROZIP1. **(C)** Apoplastic fluid from SA-treated maize leaves was co-incubated with recombinant PROZIP1. Cleavage was monitored over time by Sypro® Ruby staining in the presence or absence of E64 and/or pepstatin A (PepA). Recombinant PROZIP1 incubated in buffer alone served as control. PROZIP1 cleavage is reduced by E64 and abolished by combined E64 and pepstatin A treatment. **(D)** Apoplastic fluid from SA-treated maize was fractionated by anion-exchange chromatography to obtain a PLCP-enriched fraction containing CP1 and CP2 (Misas Villamil *et al*., 2019). Fractions were pre-incubated with pepA, EDTA and DCI, with or without E64, prior to incubation with recombinant PROZIP1. Newly generated N termini were labelled and identified by mass spectrometry and mapped onto the PROZIP1 sequence (red circles). Identified peptides are shown as grey boxes; the Zip1 region is indicated in orange and Arg-to-Ala substitutions in PROZIP1^CS^ are highlighted in red. Cleavage is restricted to Ct-PROZIP1 despite incubation with full-length PROZIP1. PLCPs and additional extracellular proteases process Ct-PROZIP1, potentially contributing to Zip1 release and peptide turnover.

We next examined whether PLCPs could process PROZIP1 after secretion. Incubation of recombinant PROZIP1 with apoplastic fluid from salicylic acid-treated maize leaves resulted in time-dependent cleavage (Fig. 4C, left), consistent with previous findings^7^. Inhibition assays showed that both the PLCP inhibitor E64 and the aspartic protease inhibitor pepstatin A^61^ partially reduced PROZIP1 cleavage when applied individually, whereas combined treatment completely abolished proteolysis (Fig. 4C, middle and right), indicating contributions from aspartic proteases and PLCPs.

To define extracellular cleavage sites, PROZIP1 was incubated with a maize apoplastic fraction enriched in active PLCPs, predominantly CP1 and CP2^62^, followed by ATOMS analysis in the presence or absence of E64 and additional protease inhibitors containing pepstatin A (aspartic proteases), DCI (serine proteases^63^) and EDTA (metalloproteases). Identified peptides exclusively mapped to the C-terminal region of PROZIP1, while the N-terminal part of the propeptide was not cleaved by the apoplastic proteases (Fig.4D). This pattern is consistent with both, the secretion of Ct-PROZIP1, lacking its N-terminal (Fig. 1), as well as with the observation that PLCPs are not required for the intracellular processing of PROZIP1.

Cleavage at positions flanking Zip1, with P2 residues valine and phenylalanine (FRR^79-81^ and VRR^102-104^) are consistent with PLCP substrate preferences^62,64^. These regions overlap with arginine-containing sites that are also cleaved by ZmMC9 (Fig. 3F), indicating that both intracellular and apoplastic proteases can potentially release Zip1 from Ct-PROZIP1. In addition, PLCP specific cleavage was also detected within the Zip1 sequence itself (for example at K^94^). This indicates that PLCPs and likely other apoplastic proteases also contribute to degradation of the Zip1 peptide.

Based on these collective data we conclude that intracellular PROZIP1 processing and the release of bioactive Ct-PROZIP1 occur independently of PLCP activity (Fig. 5). Apoplastic Ct-PROZIP1 then undergoes further proteolytic processing in the apoplast by PLCPs, likely together with additional extracellular proteases. The apoplastic cleavage can contribute to Zip1 release from Ct-PROZIP1, but it also leads to degradation of the active peptide (Fig. 5).

**Figure 5.**
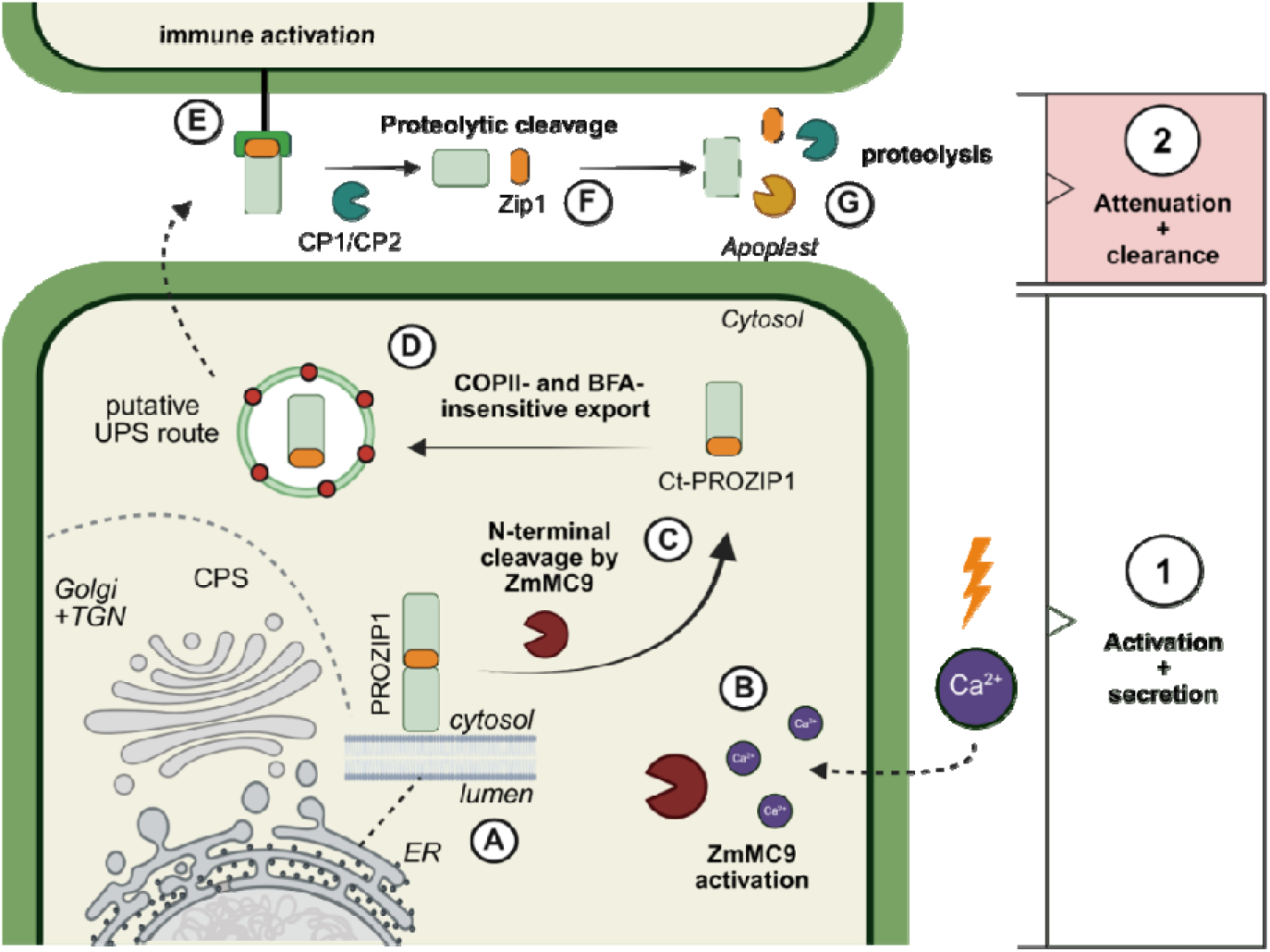
Two-step maturation of PROZIP1 generates a regulated phytocytokine signal. Schematic model of PROZIP1 processing, release and perception. **1. Intracellular activation and secretion:** PROZIP1 associates with the endoplasmic reticulum (ER) membrane after translation **(A)**. Activation of the Ca²⁺-dependent metacaspase ZmMC9 **(B)** enables N-terminal processing of PROZIP1 **(C)**, permitting release of the C-terminal fragment (Ct-PROZIP1). Ct-PROZIP1 is translocated to the apoplast via a COPII- and BFA-insensitive, independent of the Golgi and trans-Golgi network, likely involving a putative UPS route **(D)**. Ct-PROZIP1 is perceived extracellularly through N-terminal recognition of the Zip1 peptide by a yet-unidentified phytocytokine receptor, triggering immune activation in neighboring cells **(E)**. **2. Signal attenuation and clearance:** Accumulation of Ct-PROZIP1 in the apoplast promotes its cleavage by papain-like cysteine proteases (PLCPs), including CP1 and CP2 **(F)**. Excess Zip1 and Ct-PROZIP1 are degraded by additional apoplastic proteases, restricting signaling in a time- and concentration-dependent manner **(G)**.

## Discussion

Phytocytokine activity has long been assumed to rely primarily on proteolytic release, yet how intracellular activation is coordinated with extracellular signal restriction has remained unresolved. Here we define a compartmentalized two-step control mechanism in which metacaspase-dependent intracellular processing licenses the maize phytocytokine precursor PROZIP1 for secretion as a bioactive C-terminal fragment, whereas subsequent apoplastic proteolysis predominantly limits signal persistence. This spatial separation uncouples activation from attenuation and establishes proteolytic compartmentalization as a central organizing principle of Zip1 signaling.

Central to this mechanism is the type II metacaspase ZmMC9, which cleaves PROZIP1 at defined arginine residues to generate the secretion-competent Ct-PROZIP1 fragment. The dominance of the R^84^ cleavage site and the impaired processing of arginine-mutated variants indicate that substrate accessibility, rather than strict dibasic motif composition, governs productive activation. Metacaspases preferentially cleave substrates after basic P1 residues^39^, and productive cleavage at monobasic arginine sites has been demonstrated for the GRIM REAPER precursor as an AtMC9 substrate^11,40^ as well as for PROPEP1 processing by AtMC4^36^. The arginine-dependent yet motif-diverse cleavage sites identified in PROZIP1 are consistent with this substrate logic and further supported by structural modeling, suggesting that exposure of P1 arginine residues is a primary determinant of cleavage efficiency. Thus, intracellular processing of PROZIP1 reflects canonical metacaspase specificity while revealing a defined functional output: generation of a secretion-competent signaling intermediate.

*ZmMC9* is expressed alongside *Prozip1* under basal and induced conditions. As Ca²⁺-dependent activation is a defining feature of type II metacaspases^36,65–69^, ZmMC9 is likely activated by stress-associated Ca^2+^ influx, positioning metacaspase activation upstream of phytocytokine release. Intracellular metacaspase-dependent maturation has previously been described for PROPEP1 via AtMC4 and for GRI via AtMC9^11,36^, underscoring a conserved role of type II metacaspases in phytocytokine processing. However, these studies primarily demonstrated peptide release and activation without resolving how intracellular processing is integrated with extracellular signal control. In contrast, our data indicate that PROZIP1 processing defines a dominant bioactive C-terminal fragment that is secreted prior to further proteolysis and that extracellular cleavage predominantly restricts rather than activates signaling. Thus, while intracellular cleavage per se is not unprecedented, the demonstration of a secretion-competent intermediate whose activity exceeds that of the mature peptide extends existing models by revealing a spatially layered regulatory architecture.

Subcellular localization and fractionation show that PROZIP1 associates with the ER, where processed fragments accumulate in microsomal but not nuclear fractions. The insensitivity of Ct-PROZIP1 secretion to brefeldin A and Sar1 suggests that ER association does not reflect conventional ER-Golgi trafficking. Instead, ER proximity may provide a defined intracellular context for proteolytic activation prior to secretion. Similarly, non-canonical secretion routes have been proposed for PROPEP1 and PROSYS^4,70^, potentially involving brefeldin A-insensitive pathways such as EXPO-like structures^71^. Although the precise trafficking route remains to be resolved, these observations are consistent with a model in which intracellular processing constitutes a regulatory checkpoint that maintains PROZIP1 in a signaling-incompetent state until metacaspase activation licenses its release.

Once in the apoplast, Ct-PROZIP1 undergoes further proteolytic processing predominantly by PLCPs, including CP1 and CP2, which are induced during salicylic acid signaling^72^. ATOMS analyses revealed cleavage both flanking and within the Zip1 sequence. Cleavage at flanking sites was consistent with the preference of PLCPs for hydrophobic residues at the P2 position^62,64^. In contrast, cleavage within the Zip1 sequence, including at K^94^, persisted in the presence of E64, indicating that additional extracellular proteases contribute to Ct-PROZIP1 processing. Such internal cleavage events are consistent with peptide degradation rather than productive maturation. Together with the stronger biological activity of Ct-PROZIP1 compared with free Zip1, these findings indicate that extracellular proteolysis is not required for activation but instead primarily functions in signal restriction and clearance. The persistence of cleavage under PLCP inhibition further supports the view of the apoplast as a proteolytic environment that constrains phytocytokine persistence. This concept is increasingly emerging as a general principle of peptide signaling in plants, where activation mechanisms are coupled to antagonistic or degradative processes that limit signal duration. This regulatory architecture has important implications for immune control in maize. Zip1 enhances salicylic acid-associated immunity and promotes resistance to the biotrophic fungus *Ustilago maydis*^7,41^. Biotrophic pathogens depend on maintaining host viability, and excessive or prolonged salicylic acid signaling can incur fitness costs. By restricting activation to an intracellular licensing step while subjecting the exported fragment to proteolytic attenuation in the apoplast, the plant may achieve rapid amplification of defense without risking sustained or systemic overactivation. This need to fine-tune peptide-mediated immunity is further illustrated by regulatory mechanisms controlling the wound peptide systemin. The recently described anti-systemin peptide (antiSYS) is constitutively expressed and competes with a regulated systemin for binding to the SYR1 receptor, thereby establishing a basal threshold that restrains excessive defense activation^73^. Similarly, antagonistic systemin receptors with different ligand affinities (SYR1 and SYR2) modulate signaling strength in a ligand-concentration-dependent manner, providing an additional layer of immune homeostasis for the same phytocytokine signal^74^. Similarly, in PROZIP1/Zip1 signaling, the earlier transcriptional peak of *Prozip1* relative to the late detection of free Zip1^7^ supports the view that Ct-PROZIP1 represents the primary signaling entity, whereas free Zip1 detected at later time points likely reflects proteolytic processing that contributes to signal attenuation. Together, these findings highlight distinct regulatory layers including receptor antagonism, peptide competition and proteolytic attenuation that constrain phytocytokine signaling.

More broadly, compartmental separation of activation and attenuation may represent a general strategy in plant immunity. In Arabidopsis, the bacterial MAMP flg22 is both released and degraded by subtilase SBT5.2 to constrain immune signaling in space and time^75^. Similarly, CAPE9 is released extracellularly from PR1 by XCP1^27^, whereas PSK1 and related sulfated peptides require post-translational modification prior to secretion^76–78^, illustrating diverse release strategies. The PROZIP1 system differs in that activation occurs intracellularly, whereas extracellular proteases predominantly act as attenuators. Such layered control resembles regulatory principles described for animal cytokines, whose bioactivity is tightly modulated by regulated secretion, precursor processing, and extracellular inactivation^79–82^.

Taken together, our findings establish intracellular metacaspase-dependent licensing followed by extracellular restriction as a regulatory architecture that affords spatial and temporal precision to phytocytokine-mediated immune modulation. By uncoupling activation from attenuation across cellular compartments, the PROZIP1 system provides a mechanistic framework for how plants balance rapid immune amplification with controlled signal termination in a protease-rich extracellular environment.

## Material and Methods

### Plant growth conditions

Grains of maize (*Z. mays*, cv. Golden Bantam, B73 and Ky21) were placed in organic growth soil and were grown for 7 to 8 days in a plant growth chamber (15 h light at 28□°C, ∼1000□µmol□m⁻²□s⁻¹; 1 h twilight; 7 h dark at 22□°C). *N. benthamiana* plants were grown for six weeks in the university’s greenhouse at 23°C on a long day period (16 h light) and at 20°C for 8h in the dark with 30% to 40% humidity.

### Plant treatments

Zip1 peptide (QPWEGESELKLATQGASVR) was dissolved at 0.5□mg/ml. Plants were treated at 4□µM final concentration. Salicylic acid (SA) was prepared in 1% DMSO and diluted to 2□mM in water. The second leaf of 8-day-old maize seedlings was infiltrated using a 1□ml needleless syringe.

### Gene expression analyses and RNAseq

Leaf samples were harvested 6□h post-inoculation and ground in liquid nitrogen. Total RNA was extracted using TRIzol (Ambion) and treated with TURBO DNA-free (Thermo) to remove genomic DNA. cDNA was synthesized from 3□µg RNA using RevertAid H Minus and oligo(dT)₁₈ plus a *Prozip1*-specific primer. cDNA was diluted 1:100 and used for qRT-PCR with gene-specific primers and iQ SYBR Green (Promega); *Gapdh* was the reference gene. *Wrky65* was used as SA marker. RNA from Zip1-, SA-, and mock-treated samples was sequenced (Novogene) using Illumina HiSeq4000. Reads were trimmed with Trimmomatic and mapped using Bowtie2 (version 2.3.4.1)^83^. Reads were mapped to the Zm-B73 Reference-NAM-5.0 genome (maizegdb.org) and quantified using HTSeq-count (version 2.0.4)^84^.

### Microscopy and E-64 treatments of maize protoplasts

Maize protoplasts were transfected^44^ with PROZIP1-GFP, PROZIP1^CS-mut^-GFP, NLS-mCherry or mCherry-KDEL. For E64 treatment protoplasts 30µM E-64 or E-64d was added post transfection. After 16h, samples were analyzed by microscopy or Western Blot. Proteins were extracted (50mM NaOAc, pH 6), labeled for 1h with 1µM DCG04-biotin and detected using Zip1-, GFP- and streptavidin antibodies.

### *Agrobacterium*-mediated transformation of *N. benthamiana*

*Agrobacterium* strains were resuspended in 10□mM MgCl₂ (OD₆₀₀□=□1.0), mixed 1:1 with strain P19, and supplemented with 200□µM acetosyringone. Suspensions were infiltrated abaxially using needleless syringes. Leaf samples were collected 2-or 3-days post-infiltration.

### Microscopy of *N. benthamiana* leaf tissue

SP8 microscope equipped with a Plan-Apochromat 63×/1.2 NA water-immersion objective and a Galvano scanning stage (Fig. S1A, B). GFP was excited at 488□nm and detected between 500-550□nm. mCherry fluorescence was excited at 561□nm and detected between 580-630□nm. Sequential scanning was performed to avoid channel cross-talk. Additional 3D live-imaging experiments were carried out on an inverted Leica STELLARIS microscope equipped with a Plan-Apochromat 63x/1.4 NA oil-immersion CS2 objective and a Galvano scanning stage (Fig 1A, Fig. S1C). GFP was excited at 488□nm and detected between 495-515□nm. mCherry fluorescence was excited at 587□nm and detected between 605-625□nm. Prior to analysis, Leica FALCON (Fast FLIM) phasor-based separation was used to evaluate potential autofluorescence and enhance signal discrimination. No autofluorescence was detected within the specified detection windows. For 3D live imaging, infiltrated *N. benthamiana* leaves were mounted in 95% perfluorodecalin (Sigma-Aldrich, P9900-25G) in a µ-Slide 1 Well Glass Bottom #1.5H (170 µm ± 5 µm), D 263 M Schott glass (ibidi, Art. No. 82107), and covered with a coverslip to gently flatten the leaf tissue. Isolated maize protoplasts were mounted in the same type of µ-Slide and covered with a coverslip. Laser power, detector gain and offset settings were kept constant within experiments to allow comparison between constructs. Z-stacks were collected. Image processing, including brightness and contrast adjustments, was performed uniformly across the entire image using microscope software (Leica LAS X). No nonlinear adjustments were applied.

Fluorescence images (Fig. 4B) were acquired using a Nikon Eclipse Ti inverted fluorescence microscope equipped with a Plan Apo VC 60×/1.20 WI objective (Nikon). GFP and mCherry fluorescence were detected using appropriate filter sets. Images were acquired under identical exposure settings for all samples and processed using NIS-Elements software.

### Subcellular compartment separation

Transformed *N. benthamiana* plants (3□dpi) were used to isolate full extract (FE), apoplastic fluid (AF), nuclear, microsomal, and cytosolic fractions. FE was prepared by adding SDS buffer to ground leaf powder. AF was obtained by vacuum infiltration (240□mbar, 3×10□min), followed by centrifugation (2,000□g, 15□min, 4□°C). AF was fractionated using 5□kDa cut-off columns (Amicon); low and high molecular weight fractions were used for ELISA. For the high MW fraction, membranes were flushed with coating buffer. Nuclear fraction was isolated from 1.5□g FE following published protocols^85^. For cytosol and microsome we used the available protocol^86^.

### BFA A, ConA and Wm treatment and Sar1 overexpression

BFA, ConA, and Wm were infiltrated into *N. benthamiana* leaves (10□µg/mL, 0.5□µM, 35□µM) and incubated for 4□h. PROZIP1-GFP was expressed via *Agrobacterium* (mixed 1:1 with P19 ± Sar1). Apoplastic fluid was collected 2□dpi.

### GFP pull-down for cleavage site mapping

Infiltrated *N. benthamiana* leaves (3□dpi) were ground, and GFP pull-down was performed using GFP-Trap Agarose (Chromotek, gta). For maize protoplasts, 3□mL of 2×10□transfected cells were used. Pull-downs followed the manufacturer’s protocol. Ten percent of eluates were analyzed by Western blot, the rest by mass spectrometry.

### Expression and purification of recombinant PROZIP1 and PROZIP1 mutants

The PROZIP1 coding sequence was cloned into pOPINM (His-MBP tag) using the In-Fusion Dry-Down PCR Kit (Takara). Expression was performed in *E. coli* Shuffle, and protein was purified by Ni-NTA affinity and size exclusion chromatography. PROZIP1^CS-mut^ was generated following the protocol^7^. PROZIP1noR was ordered as a gene synthesis and cloned and purified following the procedure of PROZIP1.

### Structural modeling and protein-protein docking

The PROZIP1 precursor was modeled using RoseTTAFold and further refined using Refold 3^87^. The full-length amino acid sequence was used as input. The highest-confidence model based on predicted local distance difference test (pLDDT) and GDT scores was selected for further analysis.

Structural models of the maize type II metacaspase ZmMC9 were generated using AlphaFold2 Multimer^88^ implemented in localcolabfold pipeline^89^. The p20 and p10 regions were identified using Multiple sequence alignment and Interpro annotation based on the full-length ZmMC9 sequence and subsequently modeled as heterodimer to obtain mature ZmMC9. Predict ed models were inspected for structural integrity and appropriate orientation of the catalytic His-Cys dyad. All structural models were visualized and prepared for docking using PyMOL (v2.5, Schrödinger, LLC). Non-protein atoms were removed and chain identifiers were assigned.

Docking between ZmMC9 and PROZIP1 was performed using the ClusPro 2.0 protein-protein docking server (https://cluspro.bu.edu)^90^. ZmMC9 was defined as the receptor and PROZIP1 as the ligand. Docking was performed using the balanced scoring mode with default parameters and without imposing experimental restraints. Docked complexes were clustered by ClusPro based on interface similarity. The four top-ranked cluster, defined by cluster size and balanced energy score, were selected for structural inspection. Representative structures were analyzed in PyMOL to assess the spatial relationship between the predicted PROZIP1 cleavage site (R^84^) and the catalytic His-Cys dyad of ZmMC9.

### Expression and purification of recombinant ZmMC9 and ZmMC9^C141A^

The sequence of ZmMC9 (Zm00001eb150730) was retrieved from maizegdb.org. Both were amplified from 6□h Zip1-treated cDNA and cloned into a T7-based vector (C-terminal His-tag). Active site mutations were introduced following the manufacturers protocol (Agilent, Ref. 200516). Expression was performed in *E. coli* BL21.

### Activity-based protein profiling of metacaspases

Activity-based protein profiling of metacaspases was performed as described before ^56^. Recombinant protease (5□mg/mL) was incubated for 1□h at room temperature in labeling buffer (20□mM HEPES, 150□mM NaCl, 5□mM DTT) with DK13 probe (0.5□µM) and CaCl₂ (5□mM). After acetone precipitation, click chemistry was performed using a CuSO₄–TCEP–TBTA premix and Cy5. Fluorescence was detected via SDS-PAGE and Cy5 scanning.

### Metacaspase *in vitro* cleavage assays

Reactions were prepared by mixing 4× buffer (200□mM HEPES pH□7.5, 600□mM NaCl, 40% glycerol, 200□mM CaCl₂, 40□mM DTT) with water, substrate, and metacaspase in dilution buffer (25□mM HEPES, 50% glycerol). After 30□min at room temperature, reactions were stopped with Laemmli or ELISA buffer (with EDTA, leupeptin) and frozen. ELISA samples were diluted 1:10 before coating and following the published procedure ^44^.

### Proteomic Identification of protease Cleavage sites (PICS)

ZmMC9 cleavage specificity was analyzed using E. coli proteome-derived peptide libraries as described previously^57^. Purified ZmMC9 and its active site mutant (22.5□µM) were incubated with 40□µg of legumain-digested peptide library for 1□h or 16□h. Reactions were stopped with 100□µM leupeptin and 10□mM EDTA, frozen at −80□°C, acidified (0.5% TFA), and desalted using C18 STAGE tips^91^. Peptides were analyzed on an Orbitrap Elite MS (Thermo) coupled to an UltiMate 3000 RSLCnano system using a two-column setup (nanoEase™ M/Z Symmetry precolumn and HSS C18 T3 analytical column). Separation was performed at 0.3□µL/min and 40□°C with a FA-MeOH-ACN gradient. MS1 scans were acquired at 120,000 resolution; the top 15 precursors were fragmented by CID. MS data were processed with MaxQuant (v2.4.2.0), and cleavage specificity was determined with PINCIS (v12) ^57^ and visualized via the iceLogo server^92^.

### PROZIP1 cleavage site mapping

PROZIP cleavage sites were experimentally determined by Amino-Terminal Oriented Mass Spectrometry of Substrates (ATOMS) essentially as described^53^. To map PROZIP1-GFP cleavage sites *in vivo*, immunoprecipitated samples were differentially dimethyl labeled with 40□mM sodium cyanoborohydride and either light formaldehyde (^12^CH₂O; PROZIP1) or heavy formaldehyde (^13^CD₂O; PROZIP1^CS^), For *in vitro* processing with apoplast fluid, 45□µL PROZIP1 was treated with PLCP-enriched apoplastic fluid^62^ ± E-64, denatured, and labeled accordingly. All samples were split in two aliquots and digested with trypsin or legumain to increase N-termini coverage^93^. Peptides were desalted using C18 STAGE tips^91^, dried, and reconstituted in 2% ACN, 0.05% TFA. A modified ATOMS workflow with label-free quantification was applied to map ZmMC9 cleavage sites. Recombinant PROZIP1, PROZIP1^CS^ and PROZIP1^noR^ and were incubated with either recombinant active or inactive ZmMC9 or ZmMC9^C141A^ for 10 min Proteolysis was stopped by the addition of 10 mM EDTA and 500 µM leupeptin before dimethyl labeling with 40 mM formaldehyde (^13^CH₂O) and 40 mM sodium cyanoborohydrate, precipitated Sera-Mag SpeedBead magnetic carboxylate-modified particles (Cytiva) before on-bead digestion with trypsin or legumain. Peptides were eluted from the beads, dried using a SpeedVac concentrator and reconstituted in 0.1% formic acid prior to loading onto Evotips (Evotip Pure EV2013, Evosep).

Stage-tip purified peptides were separated by nanoflow HPLC (Thermo Ultimate RSLCnano equipped with Thermo Acclaim or PharmaFluidics PAC trap and analytical columns), using a 2–80% ACN gradient, and analyzed with a Bruker QqTOF (Impact II) using a CaptiveSpray ion source. MS1 scans (m/z 150–3000, 2□Hz) were followed by CID of the Top14 precursors with 45□s dynamic exclusion. Peptides loaded on Evotips in the MC9 experiment were analysed with an Evosep One system (Evosep) coupled to an Exploris 480 Orbitrap mass spectrometer (Thermo) with a Nanospray Flex ion source equipped with a stainless-steel emitter with integrated liquid junction (EV1072) mounted in an EasySpray adapter. Peptides were separated using a 21-min gradient (60 samples per day workflow) on an 8 cm × 150 µm Evosep performance column (EV1137). MS1 survey scan (m/z 350–1400) acquired at 120,000 resolution in profile mode, followed by HCD fragmentation of the 12 most intense precursors with 30s dynamic exclusion.

Acquired mass spectrometry data were analyzed with MaxQuant (v1.6.0.16, v2.0.1.0, or v2.4.20) with standard instrument settings using the *Zea mays*, improved *N. benthamiana*, or UniProt *E. coli* protein databases with appended recombinant PROZIP1 sequences and ZmMC9 sequences in the appropriate combination for each experiment. Digestion mode was set to unspecific (*in vitro*) or semi-specific (*in vivo,* MC9), with dimethylation (N-term, Lys), methionine oxidation, and N-terminal acetylation as variable modifications.

## Data availability

Source data are provided in this paper. All data supporting the findings of this study that are not directly available within the paper (and its supplementary data) will be upon reasonable request available from the corresponding authors (GD, JCMV). Raw data of RNA-seq analysis are publicly accessible in the NCBI Gene Expression Omnibus (accession number: GSE299461). Mass Spectrometry data are publicly accessible in the PRIDE database (accession numbers: PXD064635, PXD064693 and PXD075416).

## Acknowledgements

We thank Barbara Mauchbach and Sofia Dubusc for assisting with PROZIP1 construct cloning and confocal microscopy. Lukas Meschig is acknowledged for supporting cloning of ZmMC9. We are grateful to Ute Meyer for expert technical support. We thank Tolga Bozkurt and Enoch Yuen for providing Sar1A and Sar1B constructs, and to Marina Klemenčič, Katharina P. van Midden, and Renier van der Hoorn for providing the DK13 metacaspase probe and their valuable support. We gratefully acknowledge Carmen Castell and Frank van Breusegem for their valuable support in the MCA-related work. This work was supported by the Deutsche Forschungsgemeinschaft (DFG) under project DO 1421/5-2, SFB1403 (project no. 414786233), and the Cluster of Excellence on Plant Sciences (CEPLAS, EXC 2048/1—project ID: 390686111).

## Author Contributions

MK, GD, and JCMV designed the study. MK, GD and JCMV designed the experiments, PFH designed the mass spectrometry experiments. MK conducted the experiments, MK and ZS performed subcellular compartments. MK and PK performed microscopy. JCMV generated the PLCP fraction and conducted apoplastic PROZIP1 cleavage assays. BC and MS contributed to the structural modelling studies with PROZIP1. PD, MM, AP and PFH analyzed mass spectrometry data. SS provided recombinant AtMC4 and helped MK to establish MC cleavage assays. MK wrote the manuscript with contributions from all authors.

## Supplementary information

**Figure S1**. Subcellular distribution of PROZIP1 *in planta*

**Figure S2**. Arginine-dependent intracellular processing and supporting secretion analyses of PROZIP1

**Figure S3**. Arginine-dependent processing of PROZIP1 in *N. benthamiana*

**Figure S4**. Overview of type II metacaspases in relation to PROZIP1 processing

**Figure S5**. Purification and activity of ZmMC9

**Figure S6.** Purification of recombinant PROZIP1 variants and co-incubation with ZmMC9.

**Figure S7.** ZmMC9-dependent generation of Ct-PROZIP1 *in vitro*, corresponding to Fig. 4G I.

**Figure S8**. Absolute numbers of infected plants corresponding to Fig. 3G

