## Supplemental Information for "Processing and release of the maize phytocytokine Zip1"

Supplementary Figures S1 – S8

Supplementary Tables S1 – S4

### Supplementary Figures

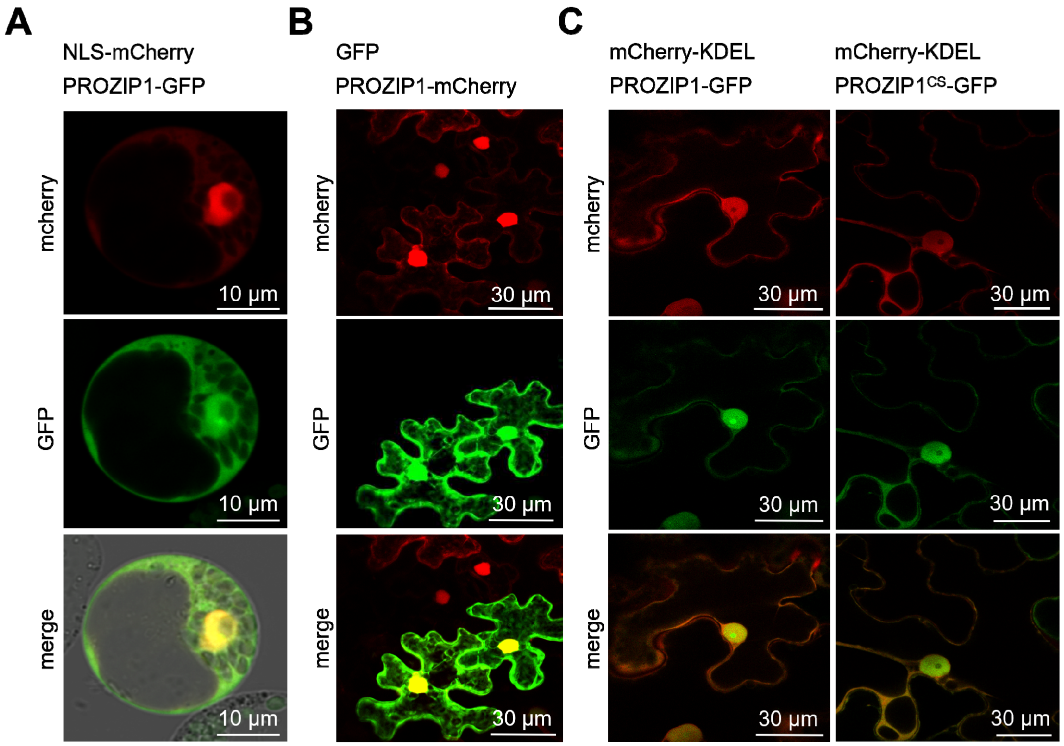


**Figure S1. Subcellular distribution of PROZIP1 *in planta*. (A)** Confocal microscopy of maize protoplasts transiently expressing PROZIP1-GFP together with nuclear NLS-mCherry. Images were acquired 16 h post-transfection. **(B)** Confocal microscopy of *N. benthamiana* epidermal cells transiently expressing PROZIP1-mCherry together with GFP. Images were acquired at 3 days post-infiltration (dpi). **(C)** Confocal microscopy of *N. benthamiana* epidermal cells transiently co-expressing PROZIP1-GFP (top) or the cleavage-site mutant PROZIP1$^{CS}$-GFP (bottom) together with the ER marker mCherry-KDEL. Shown are the mCherry channel (red), GFP channel (green), and merged images. Scale bars, 10 µm (A) and 30 µm (B, C).

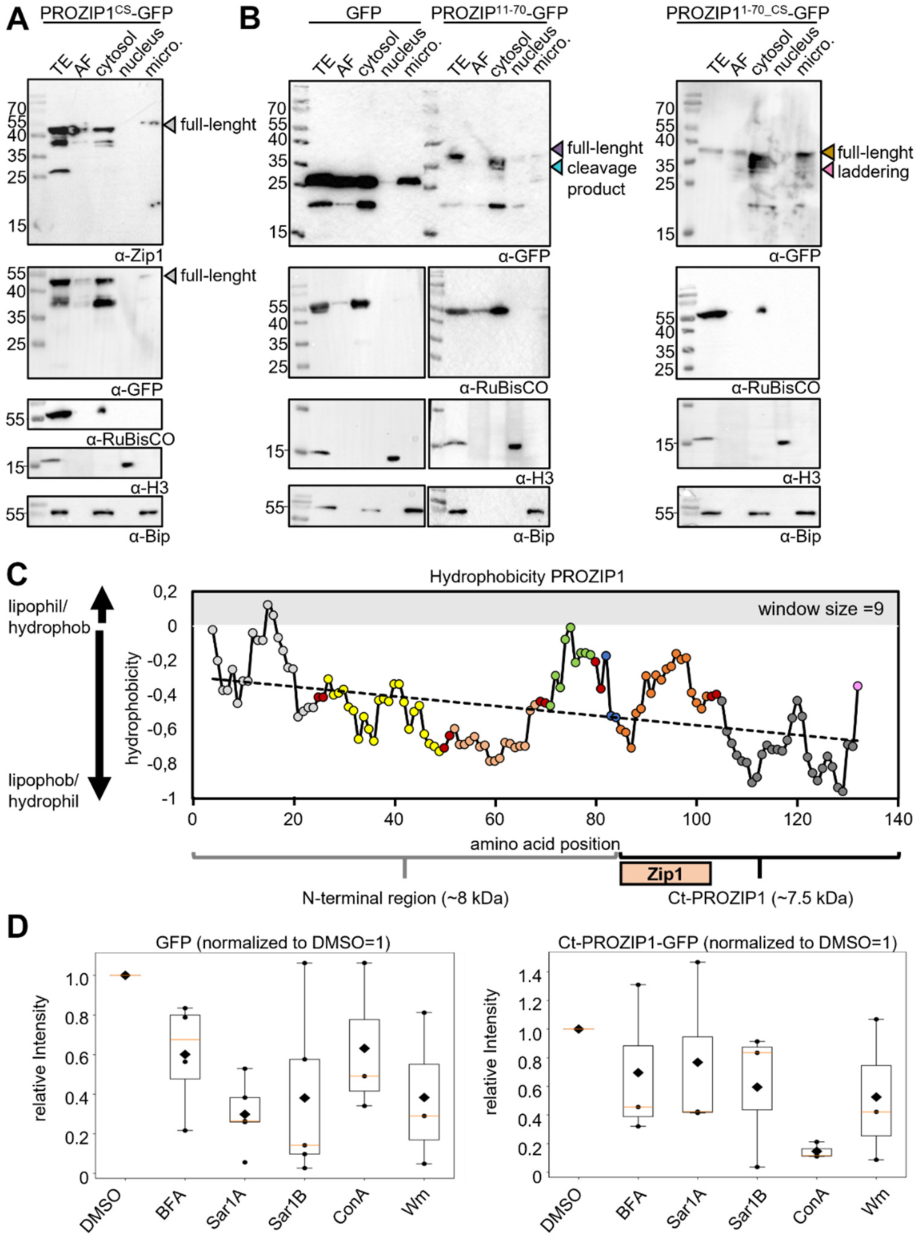


**Figure S2. Arginine-dependent intracellular processing and supporting secretion analyses of PROZIP1. (A)** Subcellular fractionation of *N. benthamiana* leaves expressing the cleavage-site mutant PROZIP1$^{CS}$-GFP. Cytosolic, nuclear and microsomal fractions, as well as apoplastic fluid, were isolated and analyzed by immunoblotting. Fraction purity was verified using RuBisCO (cytosol), histone H3 (nucleus) and BiP (microsomes). Samples were obtained 3 days post infiltration. PROZIP1$^{CS}$-GFP signals are detected in total extract (TE) apoplastic fluid (AF), cytosol, and microsomal fractions. **(B)** Immunoblot analysis of subcellular fractions from *N. benthamiana* leaves expressing PROZIP1$^{1–70}$-GFP and the RR-to-AA mutant PROZIP1$^{CS_1–70}$-GFP. PROZIP1$^{1–70}$-GFP (purple) shows discrete bands (cyan), whereas the mutant (brown) displays laddering (pink). A ~20 kDa fragment also detected in the GFP control likely reflects GFP-derived cleavage. **(C)** Hydropathy plot of the PROZIP1 amino acid sequence based on Sweet and Eisenberg (1983), showing higher hydrophobicity at the N-terminus relative to the C-terminal region. The regression curve is shown as a dashed line. Positions and sizes of the N-terminal region and the Zip1-containing C-terminal fragment (Ct-PROZIP1) are indicated. **(D)** Quantification of extracellular fluorescence signals for GFP and PROZIP1-GFP following inhibitor treatments. Values were normalized to the corresponding DMSO control of the same biological replicate (DMSO = 1). Boxplots show the distribution of five independent biological replicates; points indicate individual measurements and diamonds denote means, while the orange line indicates the median. Statistical significance was assessed using one-tailed one-sample t-tests ($P < 0.05$).

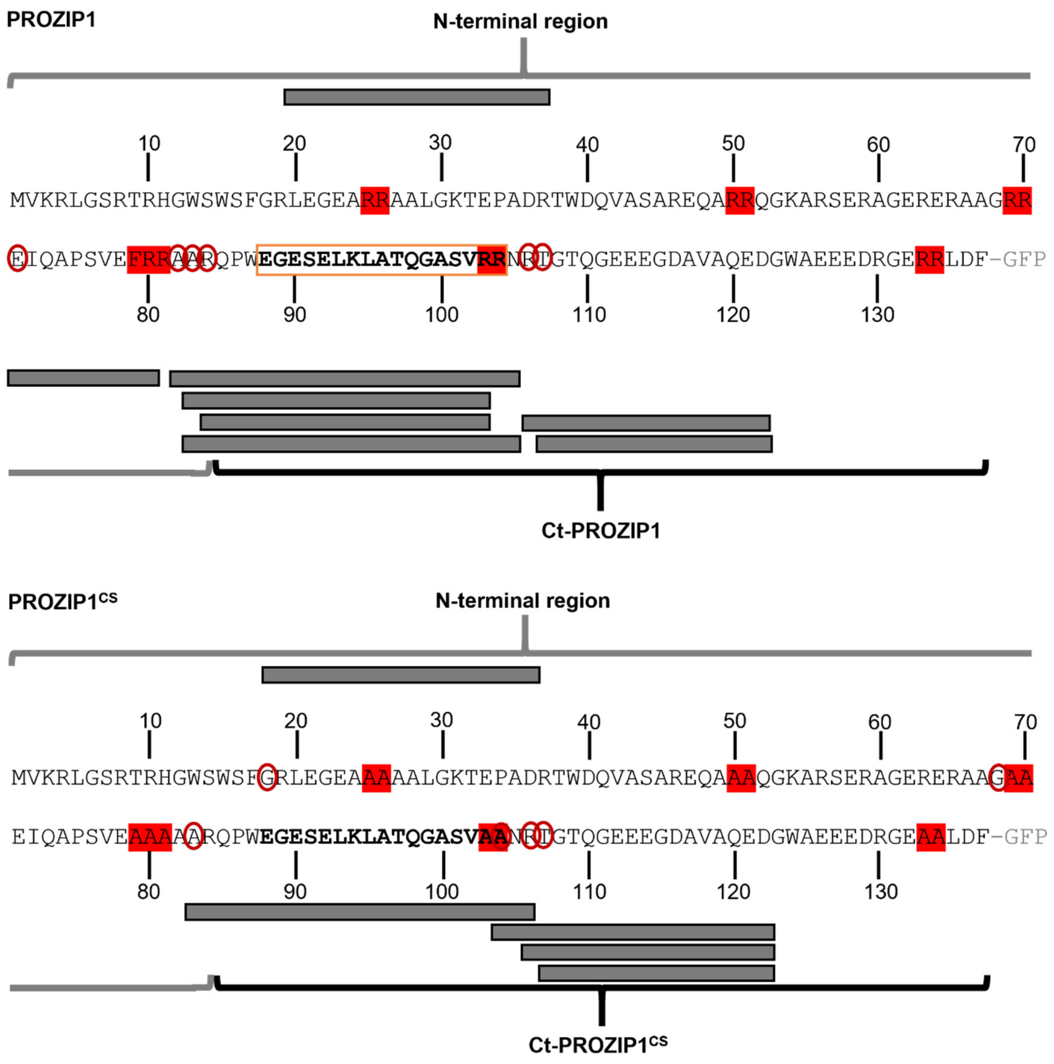


**Figure S3. Arginine-dependent processing of PROZIP1 in *N. benthamiana*.** GFP pull-downs of mCherry-PROZIP1-GFP and the cleavage-site mutant mCherry-PROZIP1$^{CS}$-GFP were subjected to N-terminal labelling by reductive dimethylation and analyzed by mass spectrometry. Identified N-terminally labelled peptides, indicating cleavage sites, are shown as grey lines mapped onto the PROZIP1 sequence. Neo N-termini are marked by red circles, and the Zip1 region is indicated by an orange box. The N-terminal region and the C-terminal PROZIP1 fragment (Ct-PROZIP1) are indicated.

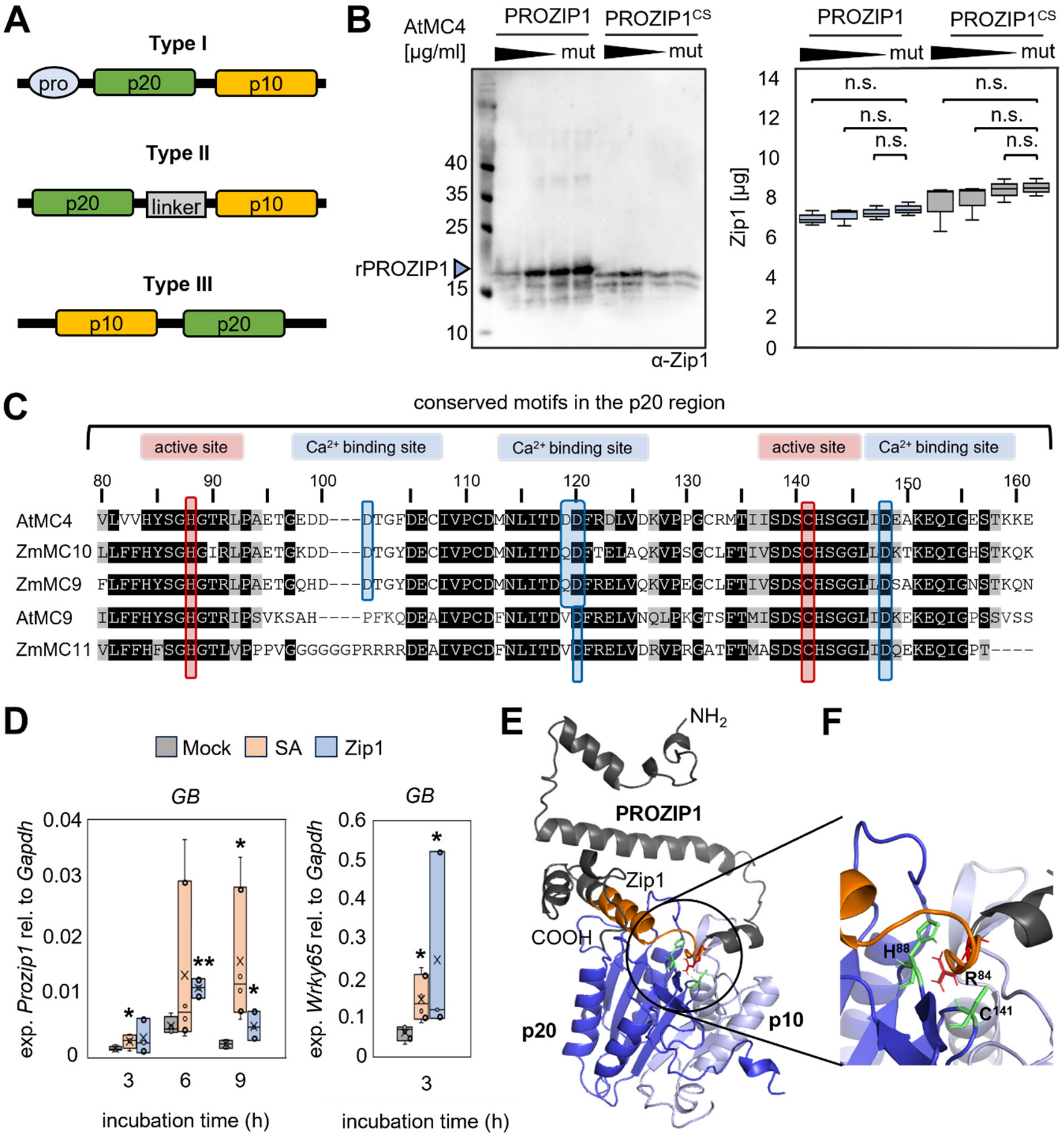


**Figure S4. Overview of type II metacaspases in relation to PROZIP1 processing. (A)** Architecture of plant metacaspases. Type I metacaspases contain an N-terminal prodomain followed by the catalytic p20 and p10 regions, type II metacaspases harbour an internal linker between p20 and p10 regions, and type III metacaspases display an inverted order of these regions. **(B)** *In vitro* cleavage of PROZIP1 by AtMC4. Recombinant PROZIP1 and PROZIP1$^{CS}$ were incubated with increasing concentrations of recombinant AtMC4 (0.2–2.5 µg ml$^{-1}$) or the inactive mutant AtMC4$^{C141A}$. Samples were analyzed by immunoblotting using an α-Zip1 antibody and by ELISA-based Zip1 detection (Koenig *et al.*, 2025). Immunoblots show concentration-dependent processing of PROZIP1 by AtMC4 but not by AtMC4$^{C141A}$, whereas ELISA signals indicate unchanged Zip1 abundance. **(C)** Multiple sequence alignment of the conserved p20 region from Arabidopsis and maize metacaspases. Conserved catalytic histidine and cysteine residues (active sites) and predicted $Ca^{2+}$-binding residues are indicated. Residue numbering corresponds to AtMC4. **(D)** Induction of *Prozip1* expression by salicylic acid (SA) and Zip1. Leaves maize seedlings (cv. *Golden Bantam*) were treated with SA, Zip1 or water (mock). Samples were collected at 3, 6 and 9 h post treatment and *Prozip1* transcript levels were quantified by qRT–PCR and normalized to *ZmGapdh*. *ZmWrky65* served as an SA-responsive marker gene (Koenig & Moser *et al.*, 2023). Data represent mean ± SEM of ≥3 biological replicates with three technical replicates; asterisks indicate significant differences relative to mock (unpaired *t*-test, $P < 0.05$). *Prozip1* is induced 3h and 9h after SA treatment and 6h and 9h after Zip1 treatment. **(E)** Structural prediction of a ZmMC9-PROZIP1 cleavage interaction. Docking model of the maize type II metacaspase ZmMC9 in complex with PROZIP1 as potential substrate. The metacaspase large (p20) and small (p10) subunits are shown in blue and light grey, respectively. PROZIP1 is shown in black, with the embedded Zip1 peptide region highlighted in orange. The predicted cleavage site $R^{84}$ is shown in red, and the catalytic $His^{88}$-$Cys^{141}$ dyad of ZmMC9 is shown in green. **(F)** Close-up view of the predicted interaction site. The docking model positions the known cleavage site $R^{84}$ of PROZIP1 near the catalytic dyad of ZmMC9.

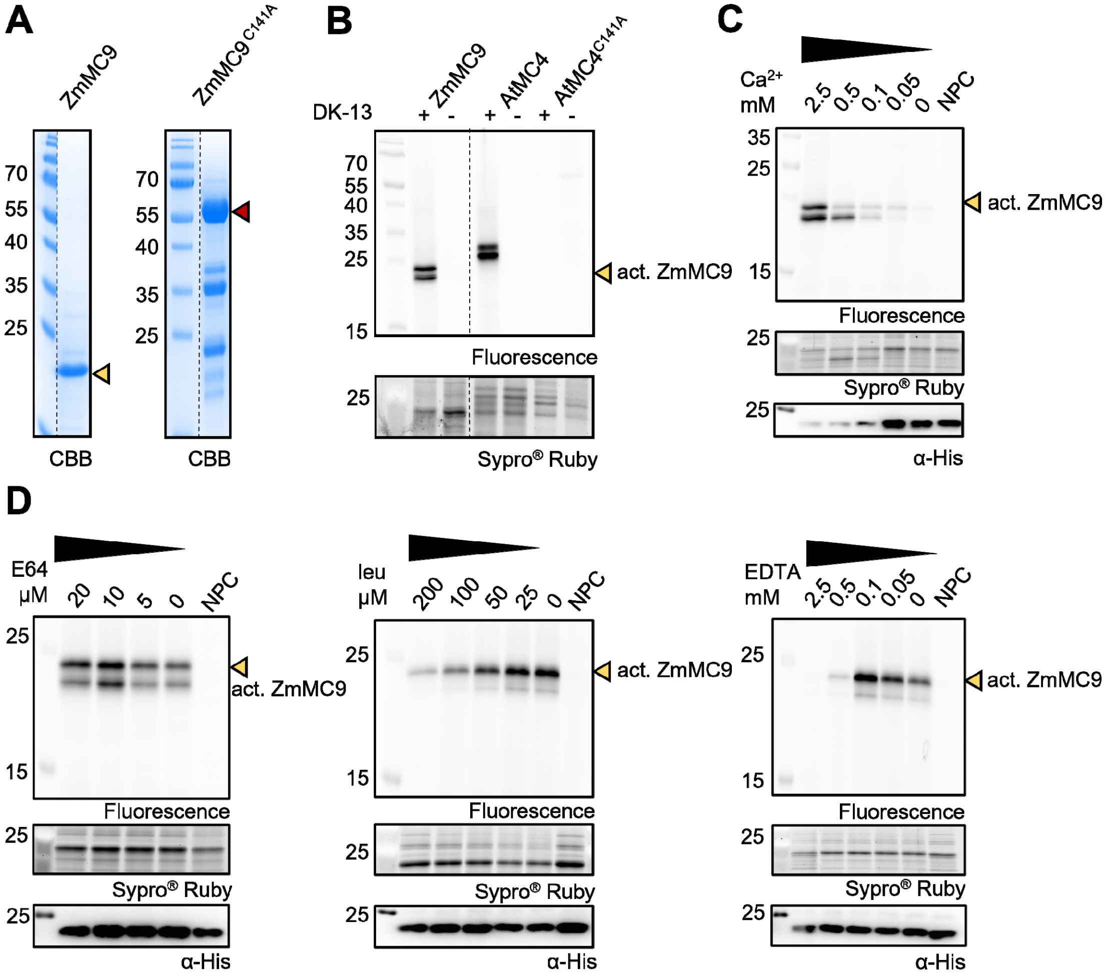


**Figure S5. Purification and activity of ZmMC9. (A)** Coomassie-stained SDS-PAGE of recombinant ZmMC9 and the inactive mutant $ZmMC9^{C141A}$. **(B)** Activity-based protein profiling (ABPP) of ZmMC9 using the fluorescent probe DK-13 (Strancar *et al.*, 2022). Recombinant Arabidopsis AtMC4 and the inactive mutant $AtMC4^{C141A}$ served as positive and negative controls, respectively. Fluorescent signals indicate active metacaspases. Sypro® Ruby staining served as loading control. **(C)** Activity-based protein profiling (ABPP) of ZmMC9 at increasing $CaCl_2$ concentrations (0 to 2.5 mM) using the probe DK-13. Equal amounts of ZmMC9 (285 µM) were used per reaction. Probe labelling was detected by fluorescent scanning. Sypro® Ruby staining and anti-His immunoblotting indicates protein loading. NPC = no-probe control. **(D)** Inhibition of ZmMC9 activity by E64, leupeptin and EDTA. ZmMC9 was pre-incubated with inhibitors prior to ABPP labelling with DK-13, followed by fluorescent detection; Sypro® Ruby staining and anti-His immunoblotting served as loading control. p20-p10 dimers (yellow) and full-length metacaspase (purple) are indicated.

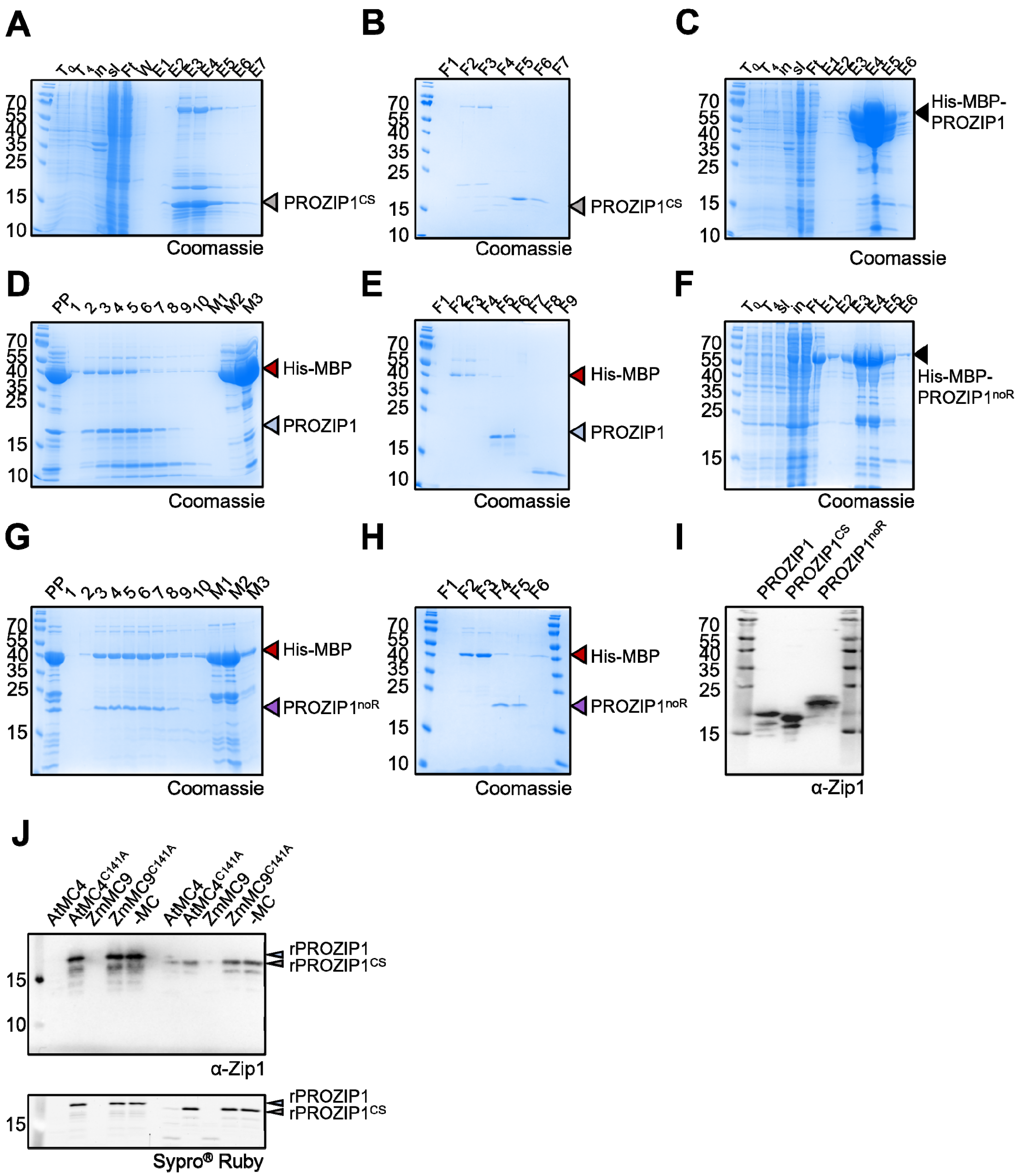


**Figure S6. Purification of recombinant PROZIP1 variants and co-incubation with ZmMC9. (A-B)** Purification of HA-PROZIP1$^{CS}$ (black) by GST affinity chromatography with on-column PreScission cleavage (A), followed by size-exclusion chromatography (SEC) (B). SEC fractions are shown by SDS-PAGE. HA–PROZIP1$^{CS}$ is indicated in grey. **(C-E)** Purification of PROZIP1 (green) by Ni-NTA affinity chromatography of His-MBP-PROZIP1 (black) (C), protease cleavage and second Ni-NTA to remove His-MBP (red) (D), followed by SEC to isolate PROZIP1 (E). **(F-H)** Purification of PROZIP1$^{noR}$ using the same strategy as in (C-E): Ni-NTA affinity purification of His-MBP-PROZIP1$^{noR}$ (black, F), protease cleavage and second Ni-NTA (G), followed by SEC (H). Lane labels: T0/4, culture before and after induction; in, insoluble; sl, soluble; Ft, flow-through; E, elution; PP, post-protease; F, SEC fractions; M, His-MBP. **(I)** Immunoblot detection of PROZIP1, PROZIP1$^{CS}$ and PROZIP1$^{noR}$ using an α-Zip1 antibody. PROZIP1$^{noR}$ migrates more slowly due to altered electrophoretic mobility caused by Arg-to-Ala substitutions. **(J)** Co-incubation of PROZIP1 (blue) and PROZIP1$^{CS}$ (orange) with ZmMC9 or the inactive mutant ZmMC9$^{C141A}$. AtMC4 and AtMC4$^{C141A}$ served as controls. PROZIP1 is cleaved in the presence of ZmMC9 and AtMC4, but remains stable in presence of the inactive variants.

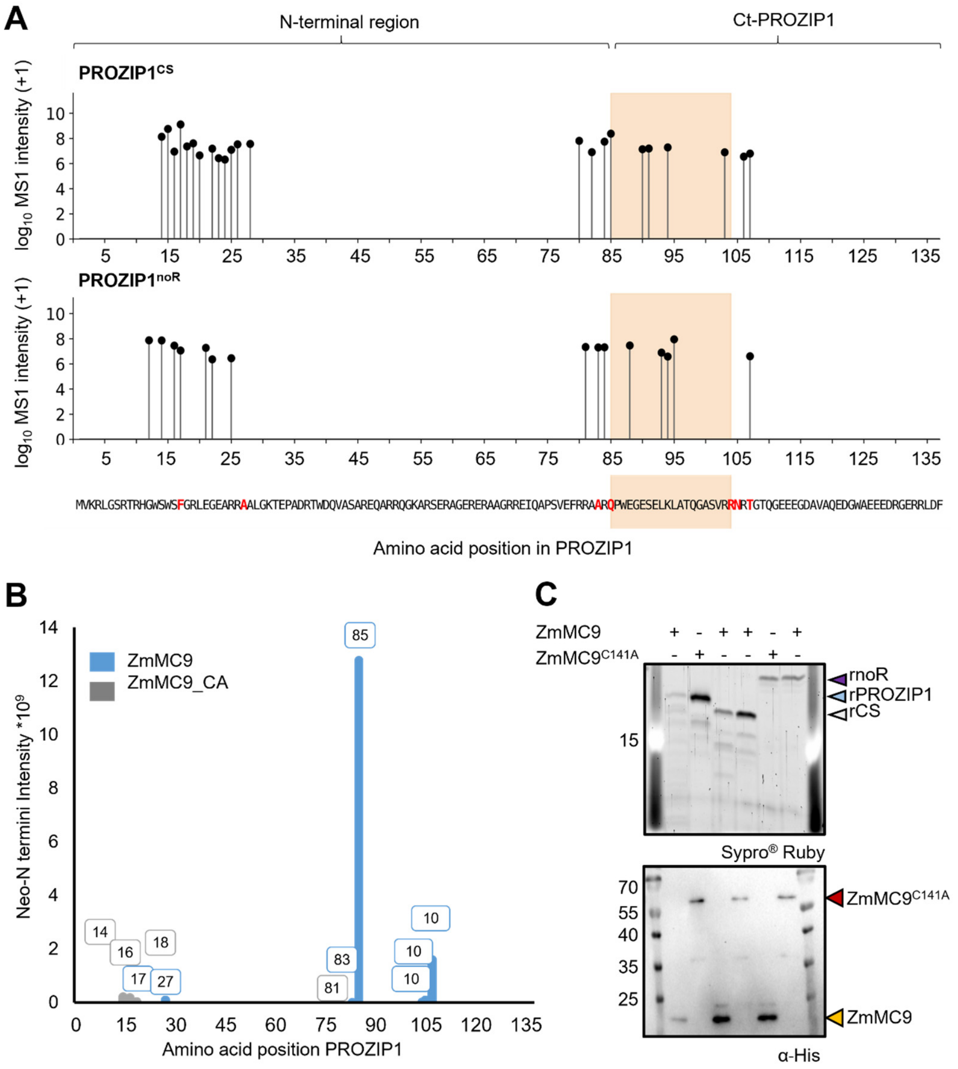


**Figure S7. ZmMC9-dependent generation of Ct-PROZIP1 *in vitro* corresponding to Fig. 4G-I.** (A) Mapping of MC9-dependent cleavage sites in PROZIP1 mutant substrates. Neo-N-termini identified by N-terminomics after co-incubation of ZmMC9 with PROZIP1$^{CS}$ and PROZIP1$^{noR}$ are mapped along the PROZIP1 sequence, using the same scale and representation as in Fig. 3F. Each lollipop indicates the position of a detected neo-N-terminus, and stem height corresponds to $log_{10}$-transformed MS1 intensity (+1) as a proxy for relative abundance. The Zip1 peptide region (amino acids 85–104) is highlighted in orange, and the PROZIP1 amino acid sequence is shown aligned to the position axis. Residues corresponding to neo-N-termini detected in wild-type PROZIP1 ($F^{17}$, $A^{27}$, $A^{83}$, $Q^{85}$, $R^{104}$, $N^{105}$ and $T^{107}$; Fig. 3F) are highlighted in red for reference. Neo N-termini were identified in the Zip1 region if ZmMC9 is co-incubated with PROZIP1$^{CS}$ or PROZIP1$^{noR}$ showing cleavage within the bioactive peptide. **(B)** N-terminomics analysis of ZmMC9-dependent cleavage. PROZIP1 was co-incubated with recombinant ZmMC9 or ZmMC9$^{C141A}$ for 10 min, followed by reductive dimethylation of neo-N termini and mass spectrometry. Identified N termini were mapped onto the PROZIP1 sequence and MS intensities are plotted (ZmMC9, blue; ZmMC9$^{C141A}$, grey), highlighting *in vitro*-generated N termini. Cleavage at $R^{84}$ which has been detected for ZmMC9 (neo N-termini $Q^{85}$) is not present with inactive ZmMC9. **(C)** *In vitro* cleavage assays. Recombinant PROZIP1, PROZIP1$^{CS}$ and PROZIP1$^{noR}$ were incubated with ZmMC9 or the inactive mutant ZmMC9$^{C141A}$ for 20 min. Cleavage was visualized by Sypro® Ruby staining. Protease loading was verified by anti-His immunoblotting. Arrowheads indicate full-length PROZIP1 (blue), PROZIP1 variants (CS=grey, noR=purple) and labelled active protease species (yellow) and its inactive full-length (red).

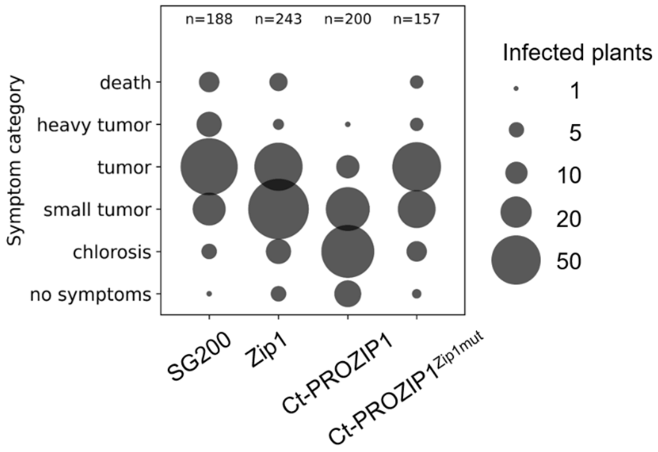


**Figure S8. Absolute numbers of infected plants corresponding to Fig. 3G.** Absolute counts of infected maize plants from the same infection experiments shown in Fig. 3G, plotted in addition to relative counts. Bubble diameter corresponds plant counts.
